# Preclinical systematic review of CCR5 antagonists as cerebroprotective and stroke recovery enhancing agents

**DOI:** 10.1101/2024.09.25.614925

**Authors:** Ayni Sharif, Matthew S. Jeffers, Dean A. Fergusson, Raj Bapuji, Stuart G. Nicholls, John Humphrey, Warren Johnston, Ed Mitchell, Mary-Ann Speirs, Laura Stronghill, Michele Vuckovic, Susan Wulf, Risa Shorr, Dar Dowlatshahi, Dale Corbett, Manoj M. Lalu

## Abstract

C-C chemokine receptor type 5 (CCR5) antagonists may improve both acute stroke outcome and long-term recovery. Despite their evaluation in ongoing clinical trials, gaps remain in the evidence supporting their use. With a panel of patients with lived experiences of stroke, we performed a systematic review of animal models of stroke that administered a CCR5 antagonist and assessed infarct size or behavioural outcomes. MEDLINE, Web of Science, and Embase were searched. Article screening and data extraction were completed in duplicate. We pooled outcomes using random effects meta-analyses. We assessed risk of bias using the Systematic Review Centre for Laboratory Animal Experimentation (SYRCLE) tool and alignment with the Stroke Treatment Academic Industry Roundtable (STAIR) and Stroke Recovery and Rehabilitation Roundtable (SRRR) recommendations. Five studies representing 10 experiments were included. CCR5 antagonists reduced infarct volume (standard mean difference −1.02; 95% confidence interval −1.58 to −0.46) when compared to stroke-only controls. Varied timing of CCR5 administration (pre- or post-stroke induction) produced similar benefit. CCR5 antagonists significantly improved 11 of 16 behavioural outcomes reported. High risk of bias was present in all studies and critical knowledge gaps in the preclinical evidence were identified using STAIR/SRRR. CCR5 antagonists demonstrate promise; however, rigorously designed preclinical studies that better align with STAIR/SRRR recommendations and downstream clinical trials are warranted. Prospective Register of Systematic Reviews (PROSPERO CRD42023393438).

## Introduction

C-C chemokine receptor type 5 (CCR5) is expressed across a variety of leukocyte subtypes, endothelial cells, and cell types in the brain (e.g., neurons, microglia, astrocytes), and is thought to play crucial roles in post-stroke neuroinflammation, blood-brain barrier repair, and neuronal survival / repair processes.^1^ CCR5 antagonists have emerged as potential therapeutic candidates for stroke, demonstrating both cerebroprotection and improved neural repair/recovery in preclinical animal models.^2–6^ However, no CCR5 antagonist drug has an approved indication in the stroke context, necessitating studies to establish safety and efficacy this population. This has led to an ongoing clinical trial to investigate efficacy of CCR5 antagonists in combination with post-stroke rehabilitation.^7^ Assessment of the preclinical evidence supporting CCR5’s role in stroke is needed to identify areas of potential benefit, and knowledge gaps, that should be addressed by future preclinical research.^8^

The stroke cerebroprotection and recovery communities have advocated for alignment of preclinical and clinical study parameters through publication of consensus recommendations for preclinical research, in an effort to enhance the translation of new stroke therapies.^9–11^ Examples include identification of more sensitive and clinically relevant preclinical outcome measures and incorporation of potentially important effect modifiers of treatment efficacy, such as age, sex, and stroke-related comorbidities (hypertension, diabetes, etc.).^9–11^ These recommendations aim to improve the translation of novel stroke therapeutics from preclinical to clinical populations, but the degree to which preclinical evidence for CCR5 antagonists satisfy these recommendations is unknown.

We sought to comprehensively evaluate the preclinical evidence for CCR5 antagonist drugs as both cerebroprotective and stroke recovery-promoting agents.^9,11^ Both perspectives are necessary to fully understand the therapeutic potential of stroke-related treatments, as distinct biological principles, time windows for treatment, and outcomes of interest underpin each of these treatment domains.^12,13^ We conducted a systematic review and meta-analysis of the CCR5 literature in conjunction with a panel of individuals with lived experiences of stroke. This review sought to explore how the preclinical evidence for CCR5 antagonist drugs aligned with guidance for preclinical stroke research provided by previous expert committees and the parameters of an ongoing clinical trial (The Canadian Maraviroc Randomized Controlled Trial To Augment Rehabilitation Outcomes After Stroke, CAMAROS, NCT04789616).^9–11^

## Results

### Study selection

Our search identified 263 citations, which was reduced to 166 unique studies after deduplication. Five studies representing 10 experiments met the eligibility criteria (Figure 1).^2–6^ Herein, “studies” refer to the published articles as a unit, while “experiments” refer to distinct investigations within each published article used to test various hypotheses (i.e., a subunit of “studies” comprised of a select cohort of animals).

**Figure 1.**
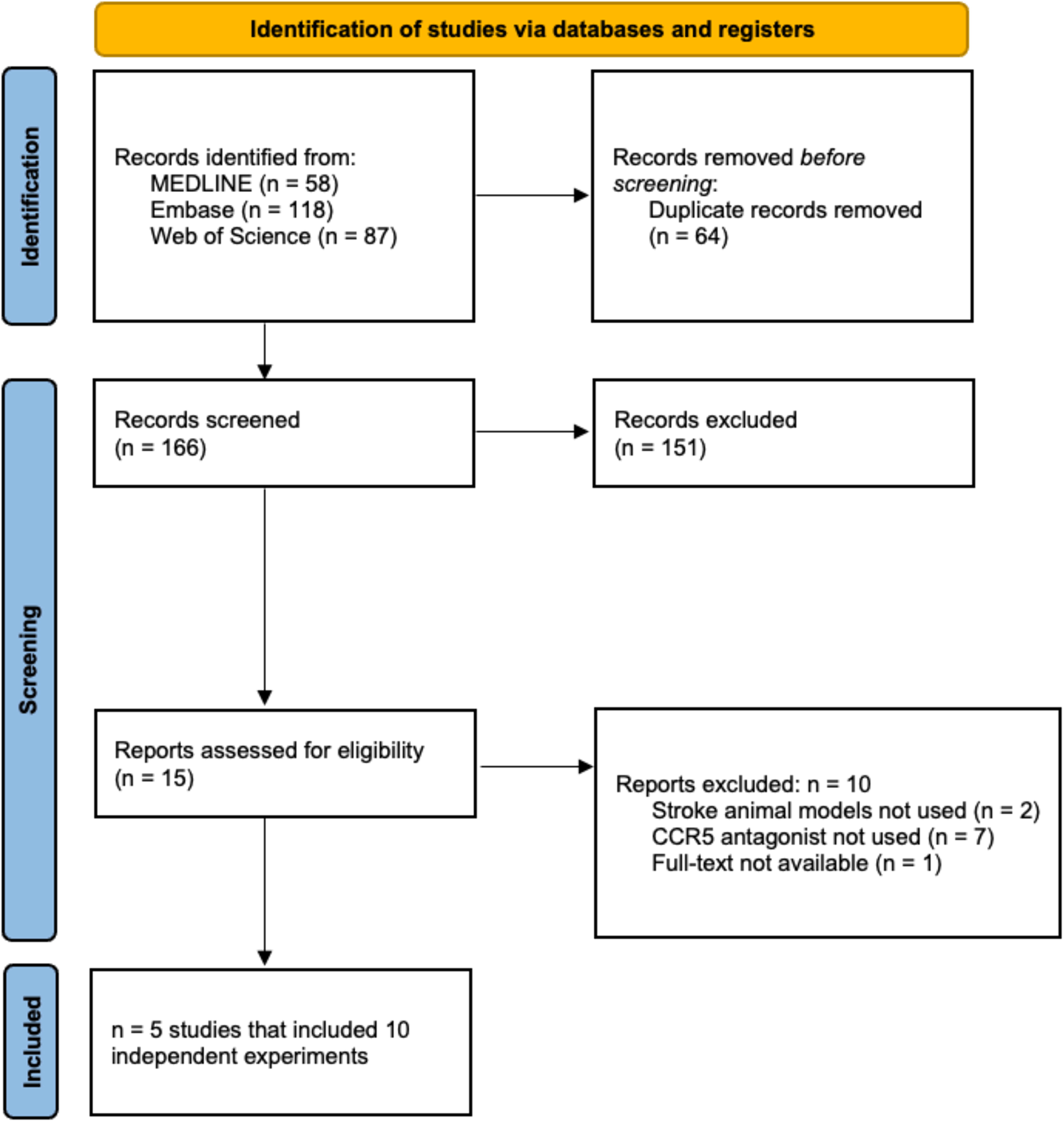
Preferred Reporting Items for Systematic Reviews and Meta-Analysis (PRISMA) flow diagram.

### Study and animal model characteristics

Most studies used ischaemic stroke (n=4/5). This was induced via intraluminal suture (n=2), cauterization (n=1), or photothrombosis techniques (n=1; Table 1). Haemorrhagic stroke was induced in one study via autologous whole blood injection. All studies used mouse (n=4) or rat (n=1) models, comprised of exclusively male animals (n=5). Relevant stroke comorbidities highlighted by patient partners and STAIR/SRRR recommendations (e.g., hypertension, diabetes) were not used in any study.

**Table 1.**
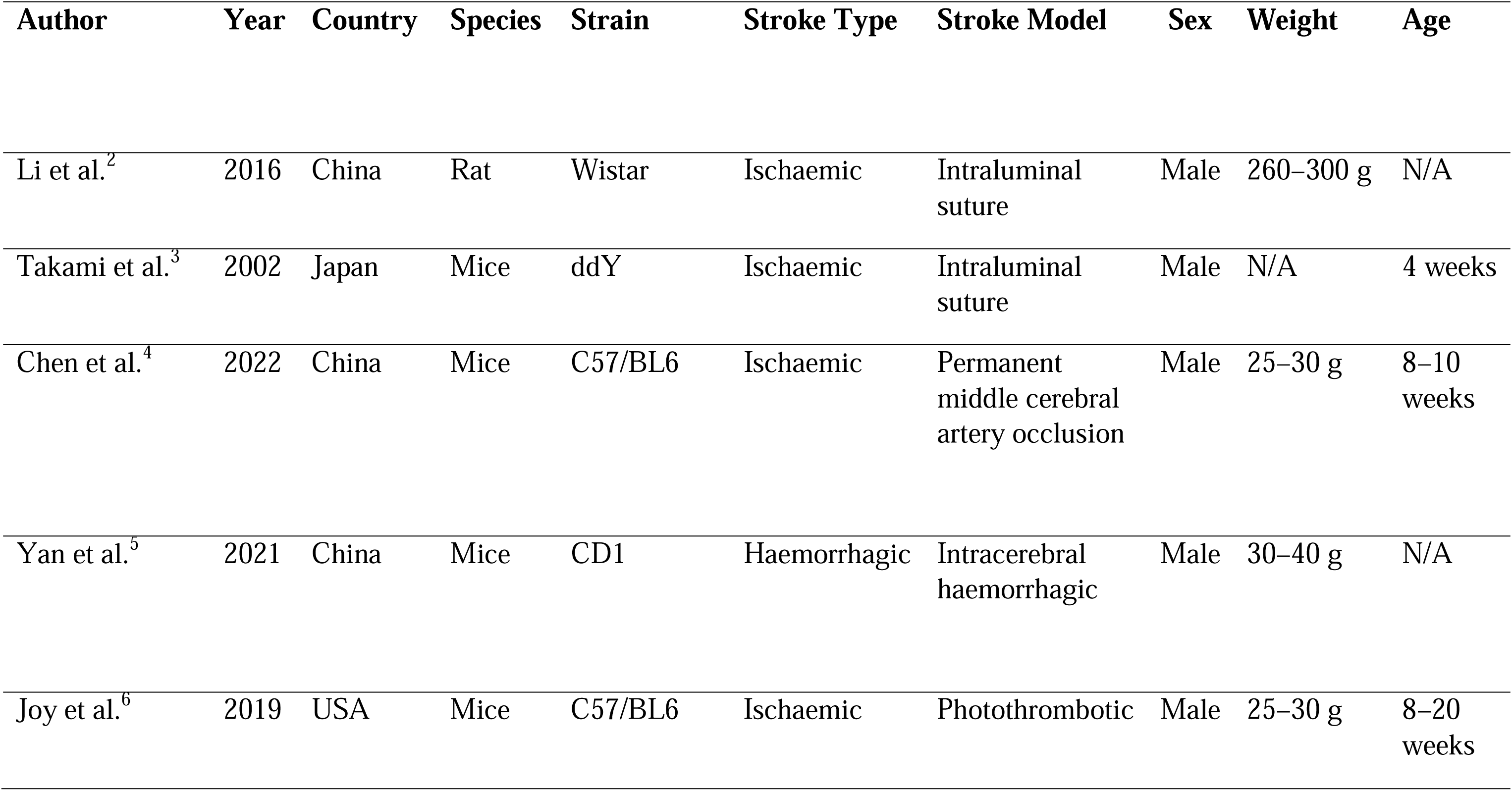
Summary of study and animal model characteristics of included articles.

| Author | Year | Country | Species | Strain | Stroke Type | Stroke Model | Sex | Weight | Age |
| --- | --- | --- | --- | --- | --- | --- | --- | --- | --- |
| Li et al.<sup>2</sup> | 2016 | China | Rat | Wistar | Ischaemic | Intraluminal suture | Male | 260–300 g | N/A |
| Takami et al.<sup>3</sup> | 2002 | Japan | Mice | ddY | Ischaemic | Intraluminal suture | Male | N/A | 4 weeks |
| Chen et al.<sup>4</sup> | 2022 | China | Mice | C57/BL6 | Ischaemic | Permanent middle cerebral artery occlusion | Male | 25–30 g | 8–10 weeks |
| Yan et al.<sup>5</sup> | 2021 | China | Mice | CD1 | Haemorrhagic | Intracerebral haemorrhagic | Male | 30–40 g | N/A |
| Joy et al.<sup>6</sup> | 2019 | USA | Mice | C57/BL6 | Ischaemic | Photothrombotic | Male | 25–30 g | 8–20 weeks |

### Intervention characteristics

Maraviroc was used in six of the experiments (n=6/10), TAK-779 in three of the experiments (n=3), and D-Ala-Peptide T-Amide (DAPTA) in one (n=1; Table 2). These CCR5 antagonists were delivered intraperitoneally (n=4), intranasally (n=2), intracerebroventricularly (n=2), subcutaneously (n=1), and intravenously (n=1) at a dose range from 0.01 to 100 mg/kg. Most studies delivered a single dose of the drug (n=6); experiments with multiple administrations (n=4) ranged from 3–63 doses. Time of initial treatment administration varied widely. Two studies (3 experiments) administered treatment pre-stroke (10-15 minutes),^2,3^ and 4 studies (6 experiments) in the acute, potentially cerebroprotective, post-stroke period (50 minutes – 24 hours post-stroke).^3–6^ One study (1 experiment) was conducted in the late sub-acute/early chronic period beginning at three to four weeks post stroke, which would be oriented towards recovery, rather than cerebroprotective, effects.^6^ Patient partners had identified several *a priori* interests (physical therapy alongside CCR5 administration, spasticity), which were also aligned with SRRR recommended considerations for preclinical stroke recovery studies.^9^ These were not reported in any included studies. We also extracted a list of outcomes used to determine CCR5’s potential mechanisms of action (Appendix 1-figure 1, Appendix 1-table 1).

**Table 2.**
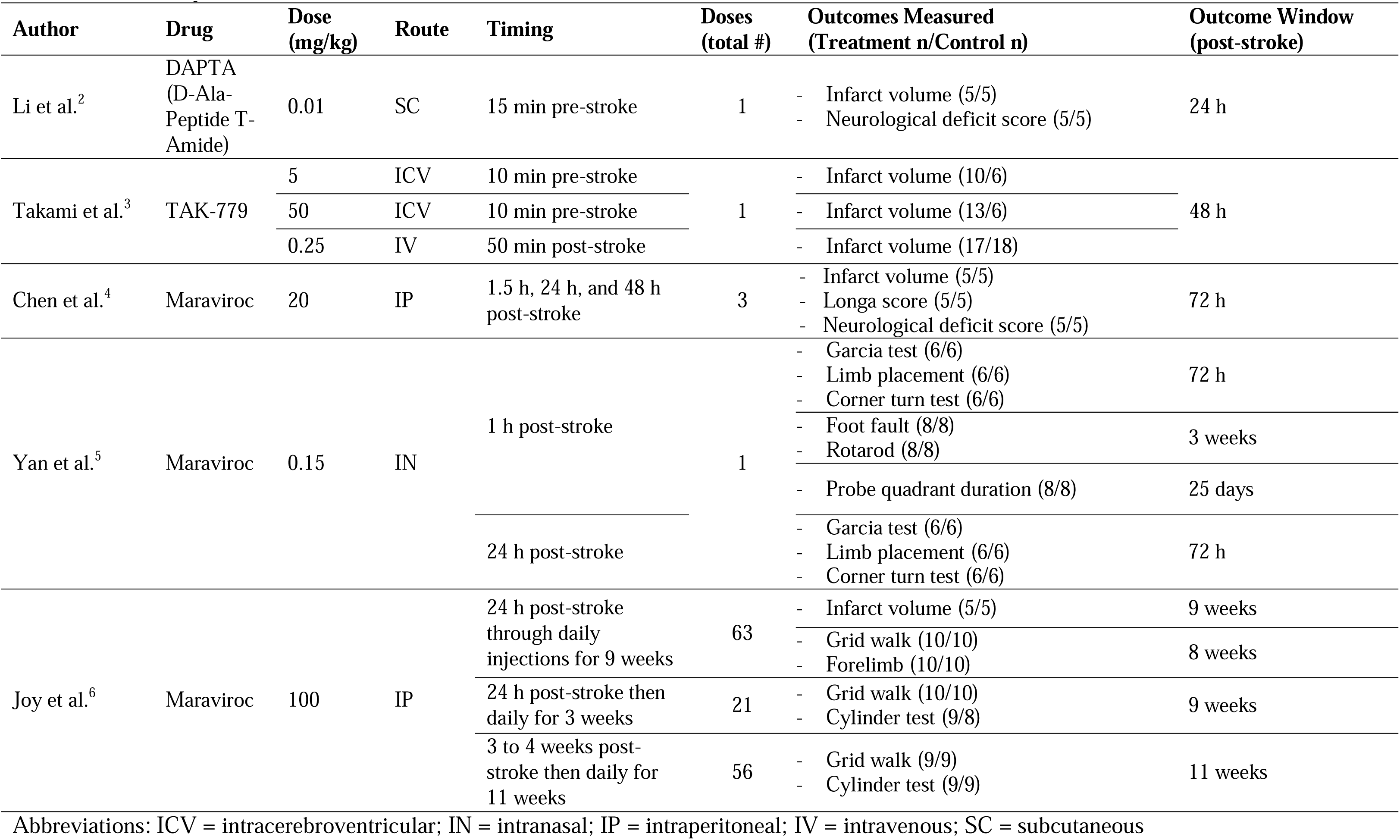
Summary of intervention characteristics.

### Meta-analysis of infarct volume

Infarct volume was reported in six experiments (n=6/10) from four different studies with an overall pooled analysis demonstrating marked cerebroprotection with CCR5 antagonists (SMD - 1.02, 95% CI −1.58 to −0.46, p < 0.0001, I^2^=34%; Figure 2). Five of these experiments measured infarct volume at 1-3 days post-stroke, and one experiment measured infarct volume at a delayed time point of 63 days post-stroke. No significant differences between pre- or post-stroke administration were observed (*P*=0.47). Post-hoc sensitivity analysis removing one experiment with extreme values^2^ demonstrated that cerebroprotection in the remaining two experiments remained statistically significant while reducing heterogeneity (SMD −0.81, 95% CI −1.25 to - 0.37, p < 0.001, I^2^=0%). A second sensitivity analysis excluded one study that measured infarct volume in mm^3^ so that all other studies could be meta-analyzed using mean differences on the percentage scale (Figure 2-figure supplement 1; MD −9.1%, 95% CI −11.6 to −6.7%, p < 0.001, I^2^=0%). This demonstrated a similar cerebroprotective effect as the other analyses. Further sub-group analyses by route of administration, time of administration, stroke model, species, CCR5 antagonist, dose, and whether behaviour tests were assessed are described in Figure 2-figure supplement 2, Figure 2-figure supplement 3, and Figure 2-figure supplement 4.

**Figure 2.**
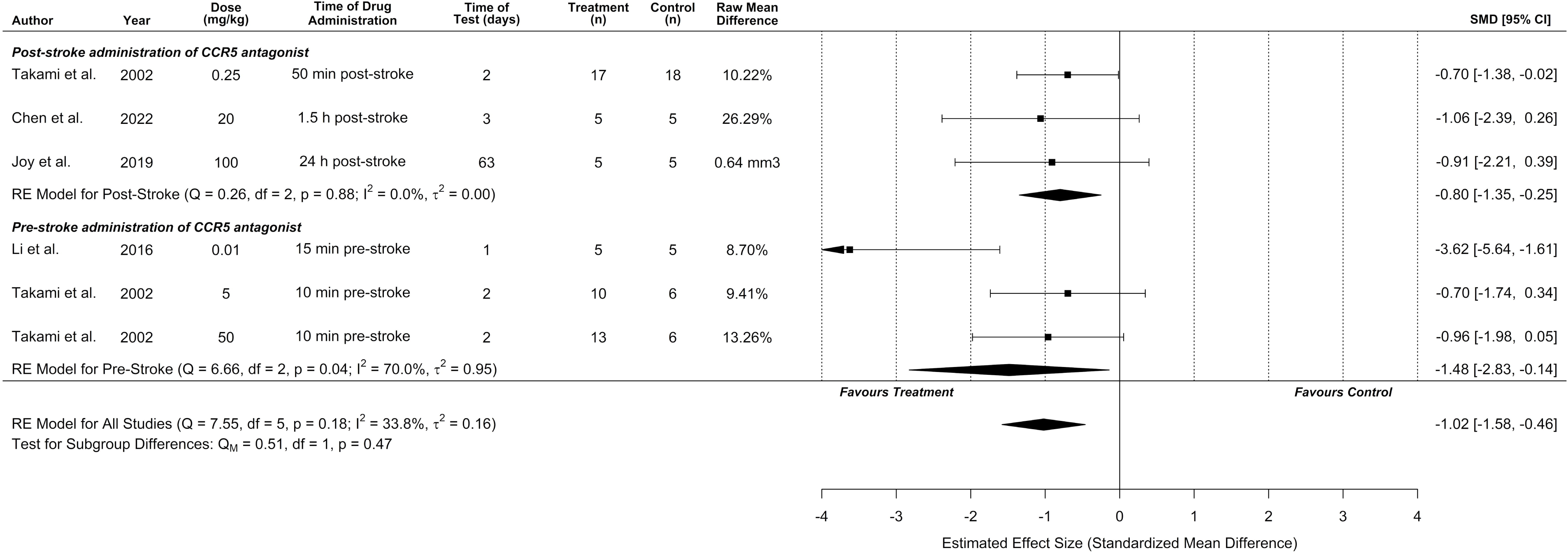
CCR5 antagonists reduce infarct volume. Data is presented as a forest plot with standardized mean differences and 95% confidence intervals. Effect sizes <0 favours drug treatment and >0 favours control. Data is stratified by timing of CCR5 antagonist administration (pre- or post-stroke induction). The ‘RE Model for All Studies’ represents a pooled estimate of the CCR5 antagonist drug effect on infarct volume from all studies combined. Separate pooled estimates are also reported for post-stroke and pre-stroke CCR5.

**Figure 2-figure supplement 1.**
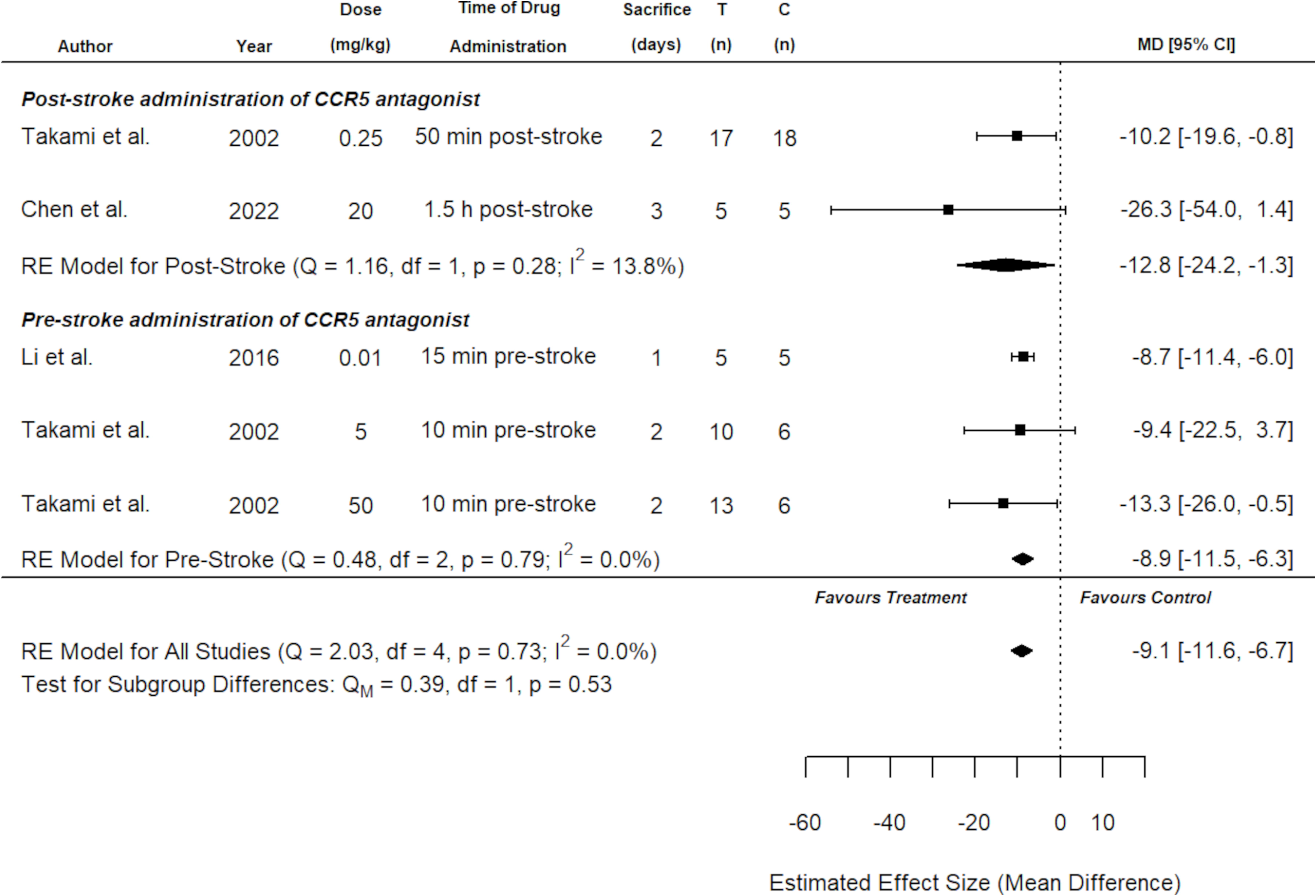
Sensitivity analysis for all included studies that reported infarct volume on a percentage scale. Overall, both the original analysis using standardized mean differences (Figure 2) and the sensitivity analysis demonstrate a significant cerebroprotective effect of CCR5 antagonists. Data is presented as a forest plot with mean differences and 95% confidence intervals. Effect sizes <0 favours CCR5 antagonist treatment and >0 favours control. Data is stratified by timing of CCR5 antagonist administration (pre- or post-stroke induction). The ‘RE Model for All Studies’ represents a pooled estimate of the CCR5 antagonist drug effect on infarct volume from all studies combined. Separate pooled estimates are also reported for post-stroke and pre-stroke CCR5. T (n) = number of animals in the CCR5 treatment group. C (n) = number of animals in the control group.

**Figure 2-figure supplement 2.**
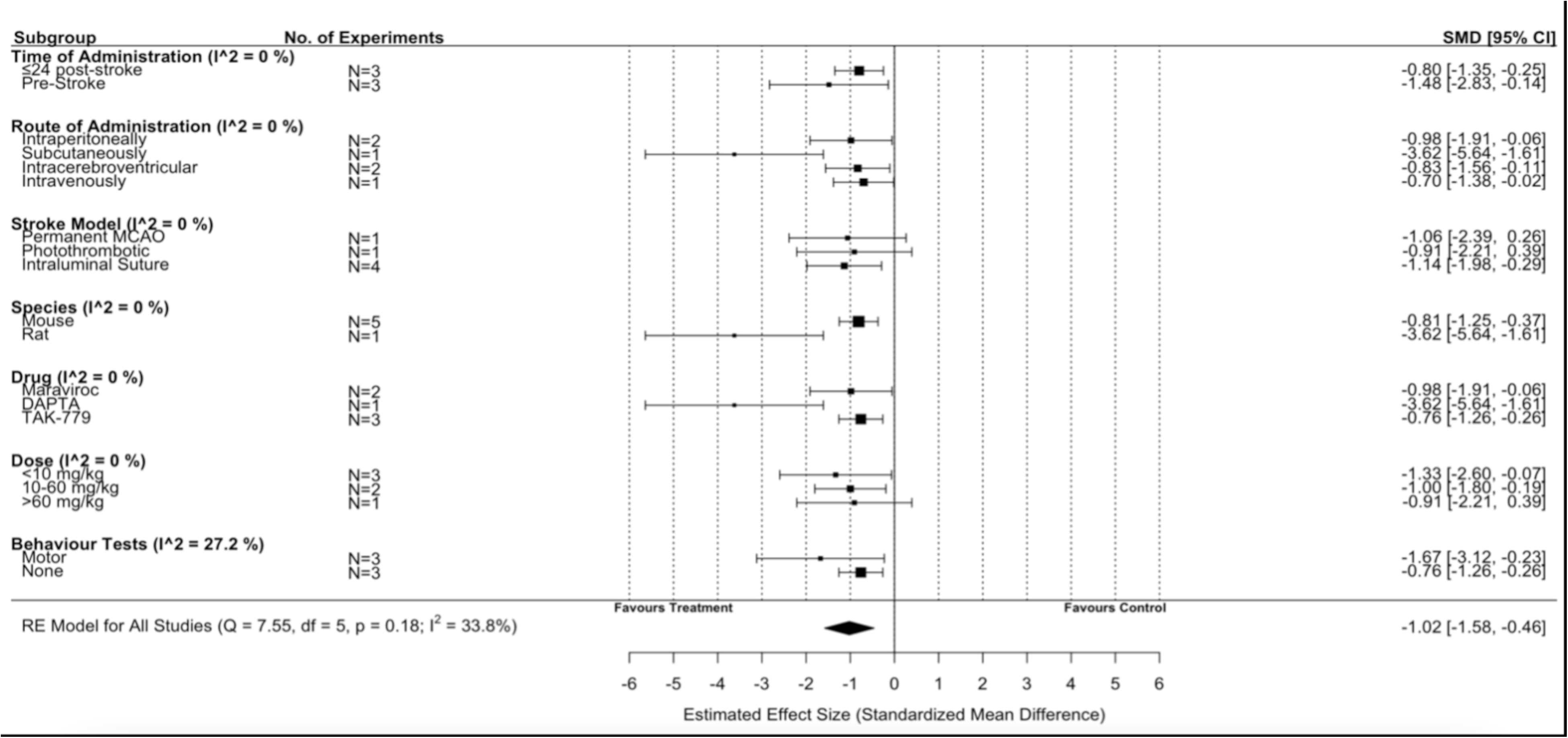
Subgroup analysis for all included studies of pre-stroke and post-stroke CCR5 antagonist administration that reported infarct volume. Each row represents pooled estimate data from studies within that subgroup. Data is presented as a forest plot with a standardized mean difference and 95% confidence intervals. The I2 value represents the statistical heterogeneity within each subgroup. Effect sizes <0 favours drug treatment and >0 favours control. The ‘RE Model for All Studies’ represents a pooled estimate of the CCR5 antagonist drug effect on infarct volume from all studies combined. Considering all six experiments (irrespective of administration timing of CCR5 antagonist), subgroup analysis demonstrated no difference in effect size when considering route of administration, stroke model, species type, drug, dose, or whether behaviour tests were assessed.

**Figure 2-figure supplement 3.**
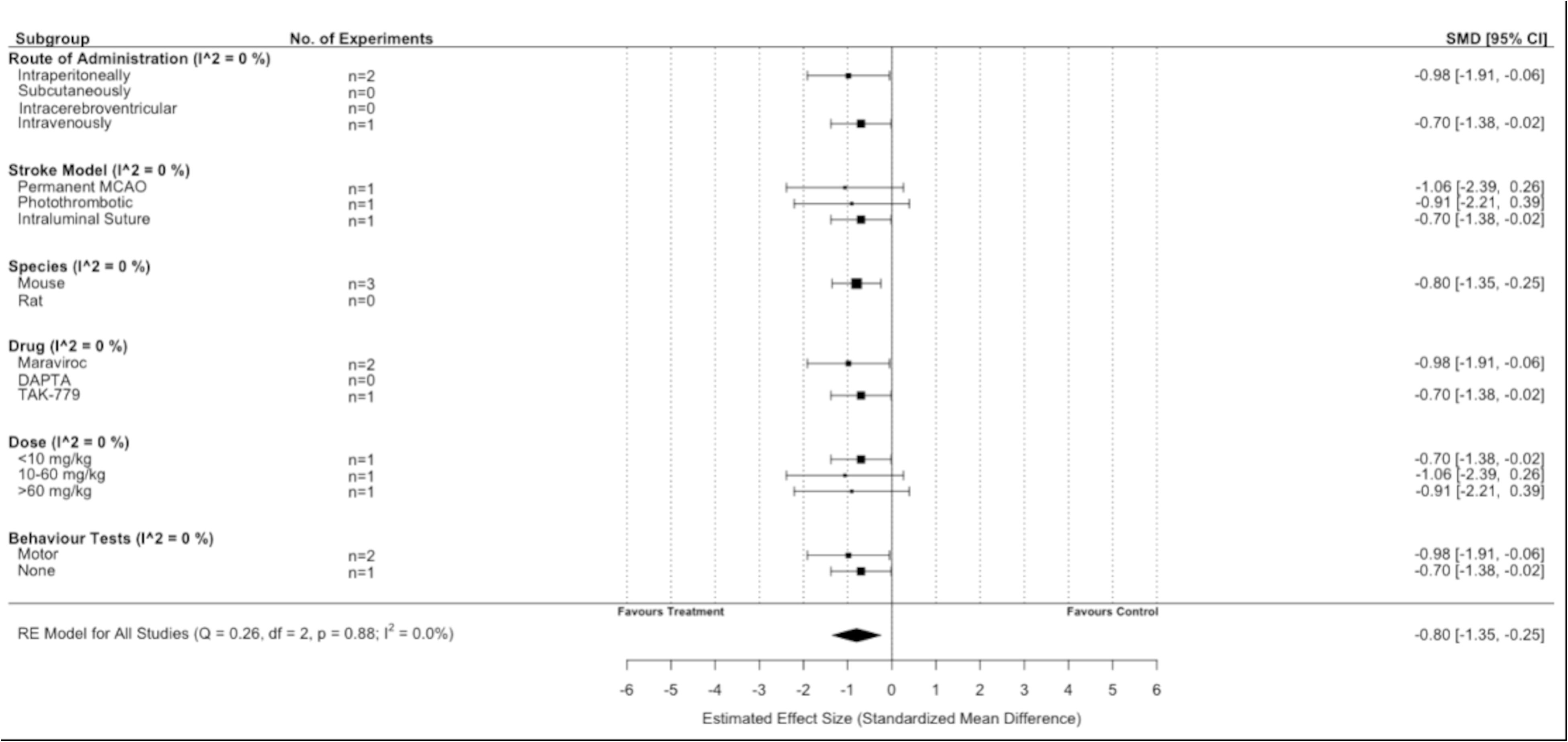
Subgroup analysis for all included studies of post-stroke drug administration of a CCR5 antagonist that reported infarct volume. Each row represents pooled estimate data from studies within that subgroup. Data is presented as a forest plot with a standardized mean difference and 95% confidence intervals. The I2 value represents the statistical heterogeneity within each subgroup. Effect sizes <0 favours drug treatment and >0 favours control. The ‘RE Model for All Studies’ represents a pooled estimate of the CCR5 antagonist drug effect on infarct volume from all post-stroke drug administration of a CCR5 antagonist studies combined. Subgroup analysis considering only studies that administered CCR5 antagonists post-stroke induction demonstrated no difference in effect size when considering any of the subgroups.

**Figure 2-figure supplement 4.**
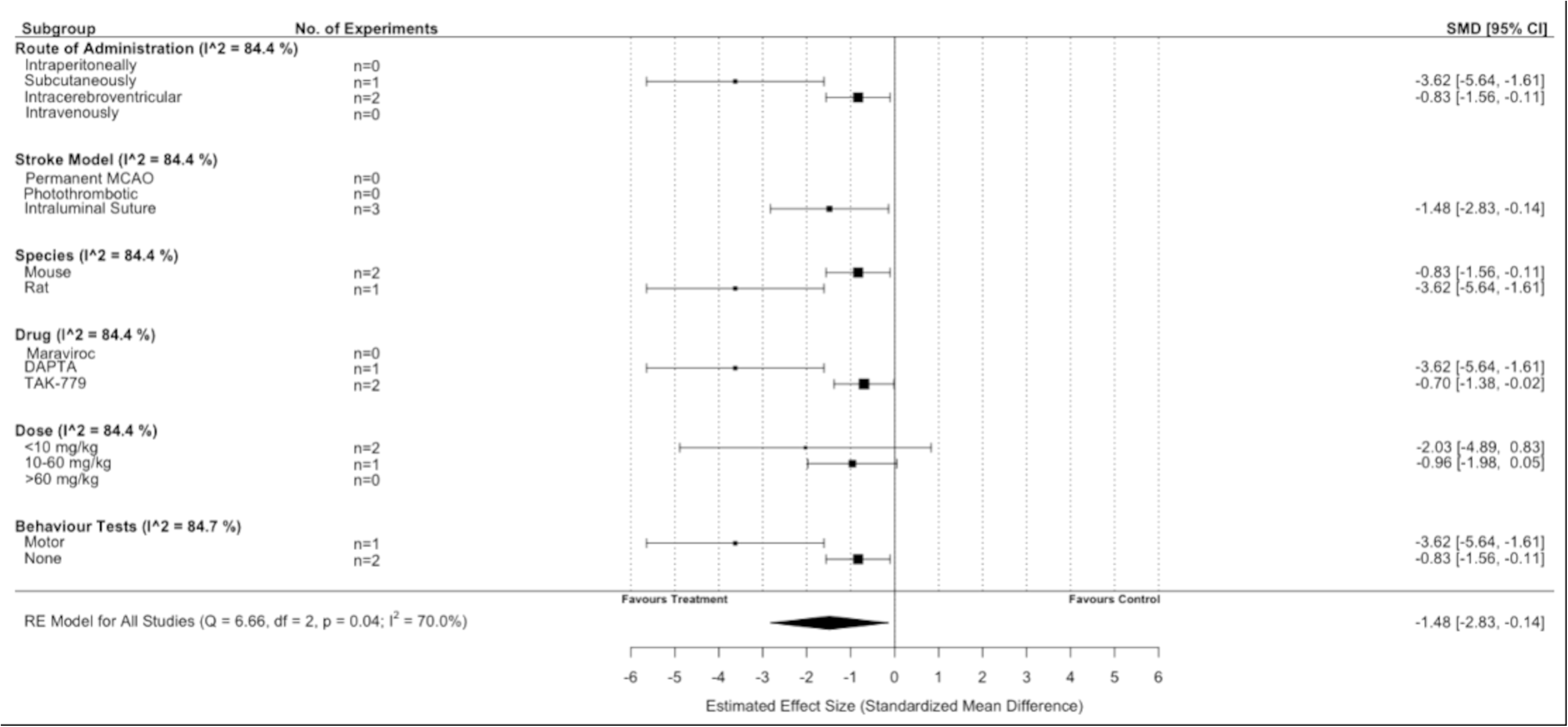
Subgroup analysis for all included studies of pre-stroke drug administration of a CCR5 antagonist that reported infarct volume. Each row represents pooled estimate data from studies within that subgroup. Data is presented as a forest plot with a standardized mean difference and 95% confidence intervals. The I2 value represents the statistical heterogeneity within each subgroup. Effect sizes <0 favours drug treatment and >0 favours control. The ‘RE Model for All Studies’ represents a pooled estimate of the CCR5 antagonist drug effect on infarct volume from all pre-stroke drug administration of a CCR5 antagonist studies combined. Subgroup analysis of studies that administered CCR5 pre-stroke induction suggested that infarct volume was reduced to a greater extent by the intraluminal suture stroke model versus other models (P=0.04), in rats versus mice (P=0.01), with DAPTA versus TAK-779 (P=0.01), and when behaviour tests were performed versus not. (P=0.01).

### Synthesis of behavioural outcomes without meta-analysis

Motor behavioural outcomes were reported in six experiments from three studies and represented seven different behavioural tasks. An additional study reported motor behavioural outcomes without standard deviations or standard errors, and thus could not be included (authors did not respond to email requests for data).^4^ A cognitive outcome (Morris Water Maze) was measured in one study. Overall, CCR5 inhibition was effective in 11 of 16 behavioural outcomes tested (Figure 3). Meta-analysis and planned subgroup analysis were contra-indicated due to an inadequate number of studies for each given outcome measure, which necessitated the synthesis without meta-analysis presented below.^14^

**Figure 3.**
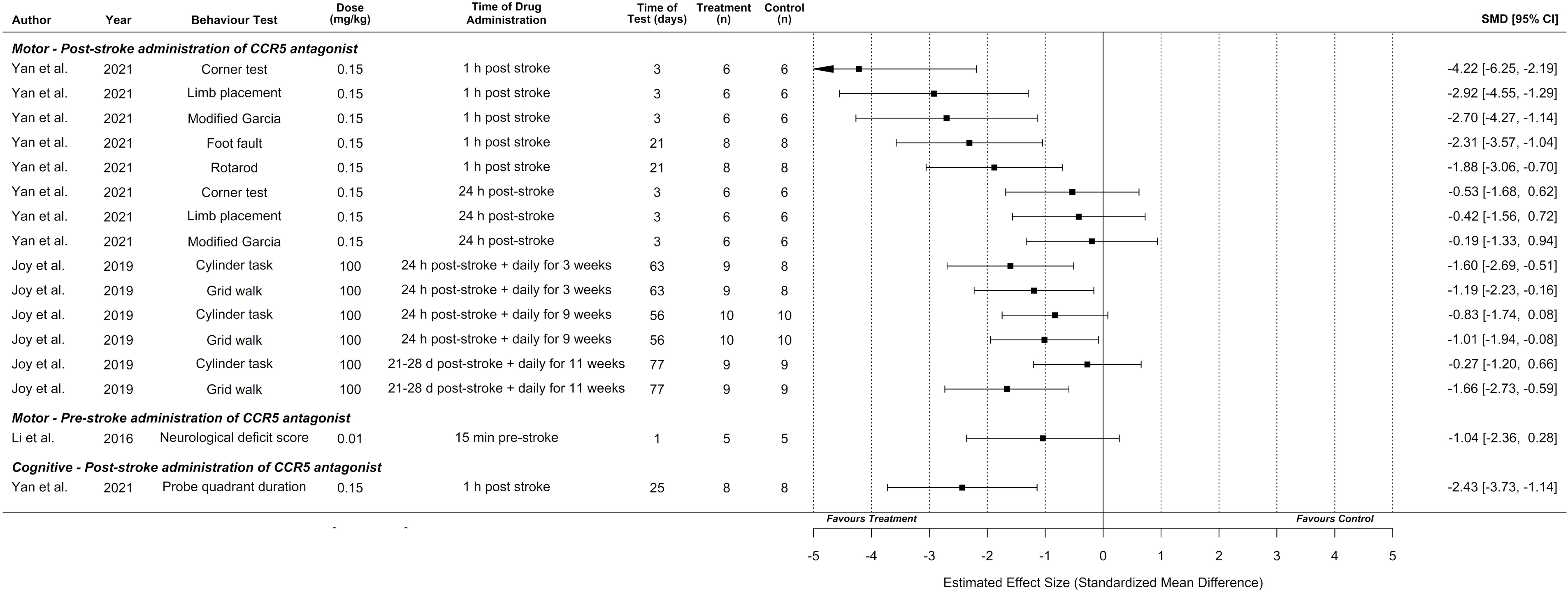
Synthesis without meta-analysis for all included preclinical CCR5 antagonist studies that reported motor and/or cognitive behavioural outcomes. Data is presented as a forest plot with a standardized mean difference and 95% confidence intervals. Effect sizes <0 favours drug treatment and >0 favours control.

Behavioural outcomes are presented by time of CCR5 antagonist administration, as discussed in the Intervention Characteristics section above, as administration time directly relates to the treatment context and patient population to which the results apply (i.e., acute cerebroprotection vs. late sub-acute/early chronic neural repair). Li et al. reported that pre-stroke administration (relevant to surgical contexts with high risk of thrombosis) of DAPTA did not result in significantly greater performance on the neurological deficit score.^2^

Regarding acute post-stroke administration times with the potential for *cerebroprotective* effects (up to 24 hours post-stroke based on observed infarct reductions in Figure 2), Yan et al. observed that CCR5 antagonist (maraviroc) administration one-hour following stroke resulted in greater motor performance on the corner test, limb placement, modified Garcia, foot fault, and rotarod tasks compared to vehicle-treated controls. Cognitive outcomes in the Morris Water Maze task (proportion of time spent in the probe quadrant) were also improved by this one-hour post-stroke administration. Significantly improved motor performance was not observed if CCR5 antagonist was administered 24 hours post-stroke, using this model of *intracerebral haemorrhage* (Figure 3).^5^ In contrast, using a *focal ischaemia* model, Joy et al. observed that CCR5 antagonist (maraviroc) administration initiated 24 hours post-stroke and continued daily for 3 weeks (which could potentially represent a mix of cerebroprotective and recovery-induced effects) resulted in greater performance on the cylinder and grid walk tasks compared to vehicle-treated controls.

Joy et al., also performed one experiment with *late sub-acute/early chronic* administration (a potentially critical time for neural repair) initiated at three to four weeks post-stroke and continued daily for 11 weeks, that demonstrated significantly improved performance for the grid walk, but not cylinder, task.^6^ This experiment was initiated outside the plausible range for a cerebroprotective effect, implying behavioural improvement involved *recovery-promoting mechanisms*. However, equivalent infarct volumes were not demonstrated between the treated and control groups in this cohort, which could potentially lead to confounding effects.

### Risk of bias

All articles had a ‘high’ risk of bias in at least one domain of the SYRCLE tool.^15^ Most domains within each study demonstrated an ‘unclear’ risk of bias. All studies reported randomizing animals; however, as commonly observed in the preclinical literature,^16–18^ only one of these studies provided sufficient detail to ensure that the randomization method had a low risk of bias. The SYRCLE domains with the highest risk of bias were incomplete outcome data, with 80% of studies (n=4/5) failing to provide complete data for all animals initially included in the study, as well as selective outcome reporting, with 60% (n=3/5; all studies of maraviroc) not providing complete data for all expected outcomes discussed in the methods (Figure 4).

**Figure 4.**
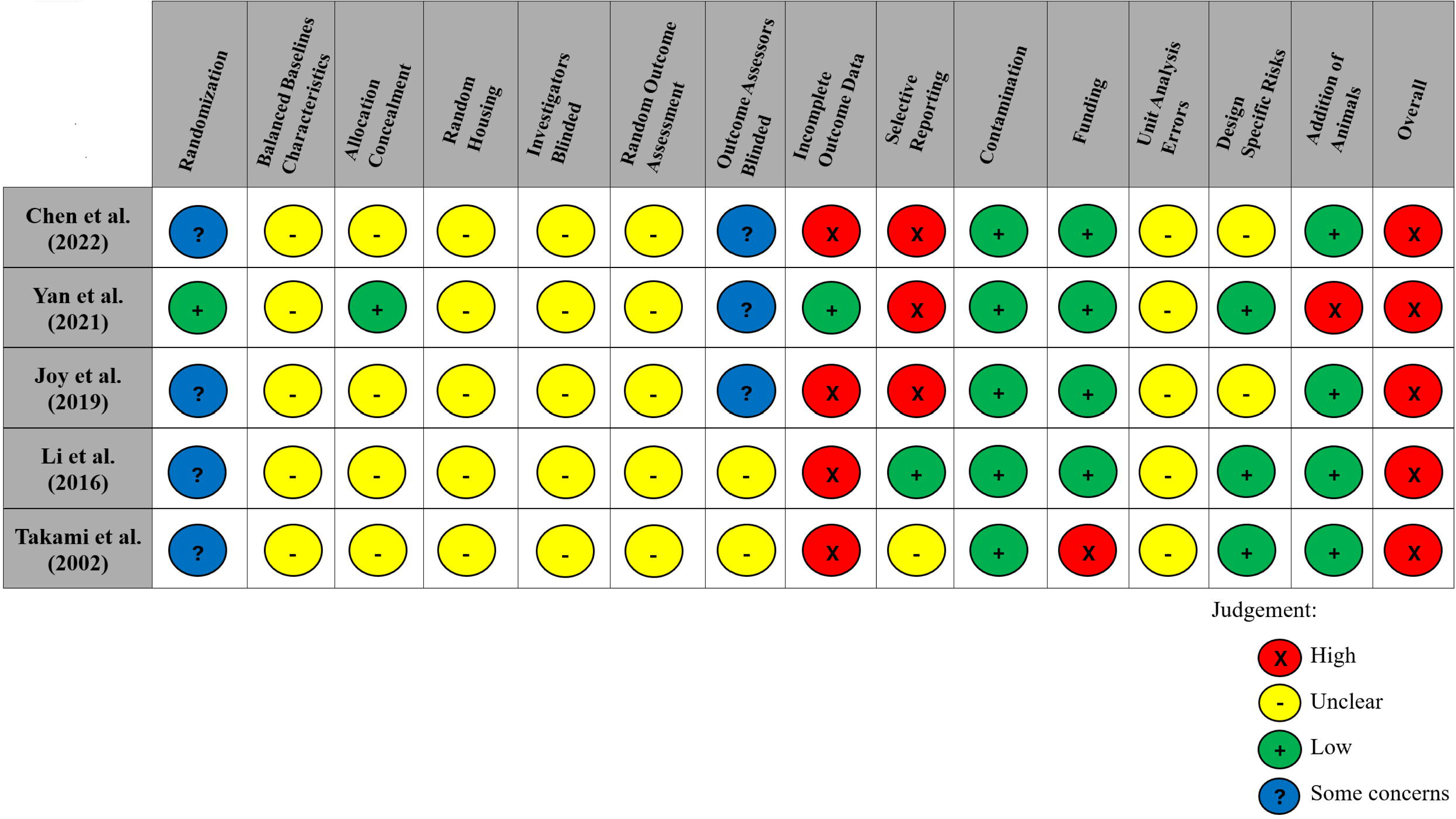
Modified risk of bias traffic light plot in accordance with the SYRCLE tool. Yellow represents an unclear risk of bias, green represents a low risk of bias, and red represents a high risk of bias. Blue represents some concerns of a risk of bias. The risk of bias was ‘unclear’ across all studies for the domains of baseline characteristics because of missing data in the studies, random housing because no details on this domain were reported, and random outcome assessment because no details of how cohorts of animals were selected to perform certain outcomes nor how the order of outcome assessment proceeded. Four studies did not report on allocation concealment, and two studies did not report on blinding investigators and outcome assessors and were deemed as having an ‘unclear’ risk of bias. 80% of studies exhibited a ‘high’ risk for incomplete outcome data. Similarly, three studies (60%) had a ‘high’ risk of selective outcome reporting since all expected outcomes discussed in the methods of the articles did not align with their results. Other potential sources of bias considered included the source of funding (industry funded), contamination of pooling drugs (additional treatment which might influence or bias the result), unit error analysis (all animals receiving the same intervention are caged together, but analysis was conducted as if every single animal was one experimental unit), design-specific risks of bias (reporting details of which animals performed the same or different outcomes), and the addition of new animals to replace dropouts from the original population. Two studies (40%) had a ‘high’ risk in at least one of these additional categories.

### Comprehensiveness of preclinical evidence and alignment with clinical trials

We assessed comprehensiveness of the preclinical evidence using the STAIR and SRRR recommendations^9,11,19,20^ as well as alignment with study parameters of the CAMAROS trial.^7^ A summary of assessment items from STAIR XI is provided in Table 3, with additional items from STAIR I, VI, and SRRR recommendations included in Table 4.

**Table 3.** Alignment of included preclinical studies with STAIR recommendations and CAMAROS trial parameters.

| Recommendation | Overall preclinical evidence | CAMAROS trial alignment | Notes |
| --- | --- | --- | --- |
| **Candidate treatment qualification** |  |  |  |
| (1) Establish treatment dose-response | Yes | No | • **Preclinical** – doses across all studies from 0.01 to 100 mg/kg<br>• 6 doses tested for infarct volume, 3 doses for behavioural effects<br>• Joy et al. demonstrated that 100 mg/kg of maraviroc in mice results in cerebrospinal fluid levels lower than in humans (13.8±5.4 vs. 33.6–60 ng/mL)<br>• **CAMAROS** – participants take 300 mg doses twice daily for 8 weeks<br>• **Interpretation** – maximum preclinical dose may not represent clinical levels |
| (2) Treatment given after clinically relevant delayed times (one to 4.5 hours post-stroke) | Yes | Yes | • **Preclinical** – 3/5 studies (all maraviroc) administered drug from 1 to 24 hours post-stroke (acute phase)<br>• Joy et al. initiated maraviroc administration at 20 days post-stroke for 11 weeks (subacute phase)<br>• **CAMAROS** – participants recruited between 5 days and 6 weeks post-stroke (acute / early subacute)<br>• **Interpretation** – one experiment in a time window relevant to clinical trial |
| (3) Both histologic and behavioural outcomes at acute (1 to 3 days) and chronic (7 to 30 days) time points | Yes | Yes | • **Preclinical** – 2/5 studies included both histologic and behavioural outcomes<br>• Mainly tasks of spontaneous movement (e.g., cylinder) or motor coordination (e.g., grid walk)<br>• Tests ranged from 3 days (acute) to 11 weeks (chronic) post-stroke for all included studies, but short term assessment may only delay cell death<sup>21</sup><br>• **CAMAROS** - motor learning (Fugl-Meyer Upper Extremity Assessment Score and 10-Meter Walk Test Score) measured as the primary outcome at baseline, 4-week (late subacute), 8-week (late subacute), and 6-month (chronic)<br>• **Interpretation** – challenging to directly compare preclinical and clinical motor tasks, but testing range is comparable. Most relevant Joy et al. experiment (subacute administration) did not measure histologic outcomes |
| (4) Treatment reaches presumed target and causes expected physiological effects that can be assessed with a clinically relevant biomarker | Yes | NA | • **Preclinical** – 5/5 studies demonstrated trends for CCR5 antagonism to reduce infarct volume. Plausible that CCR5 influences stroke-relevant mechanisms<br>• **CAMAROS** – does not include outcomes to assess treatment mechanisms |
| (5) Treatment able to pass the blood-brain barrier | Yes | NA | • **Preclinical** – 1/5 studies used mass spectrometry to demonstrate that maraviroc is present in the brain and cerebrospinal fluid<br>• **CAMAROS** – does not include outcomes to assess presence of drug in the brain |
| **Preclinical assessment / validation** |  |  |  |
| (1) Aging/adult age | No | No | • **Preclinical** – 4/5 studies used rodents with weights/ages consistent with young adulthood; 0/5 studies used rodents ≥10 months old.<br>• **CAMAROS** – Age ≥18 year old adult participants eligible for recruitment<br>• **Interpretation** – Preclinical studies consistent with CAMAROS eligibility criteria; however, no preclinical study examined middle-aged or elderly animals that would be more representative of the trial and overall stroke<br>populations |
| (2) Male and female animals | No | No | • **Preclinical** – 0/5 studies included female animals<br>• **CAMAROS** – both male and female sexes eligible for recruitment<br>• **Interpretation** – effects of CCR5 antagonists for stroke recovery in female animals represents a critical knowledge gap |
| (3) Animals with co-morbidities | No | No | • **Preclinical** – 0/5 studies included animals with common stroke co-morbidities (e.g., diabetes, hypertension, etc.)<br>• **CAMAROS** - participants with stroke co-morbidities eligible for recruitment<br>• **Interpretation** - effects of CCR5 antagonists for stroke recovery in animals with common stroke comorbidities represents a critical knowledge gap |
| (4) Evidence from two or more laboratories | Yes | No | • **Preclinical** – 5/5 studies demonstrated stroke-related benefits; however, no study had their results replicated by independent laboratories<br>• **CAMAROS** – trial protocol will be replicated by multiple study sites<br>• **Interpretation** – future preclinical experiments could consider assessing effects using a multilaboratory approach, or independently replicate the effects of existing preclinical studies |
| (5) Gyrencephalic species | No | No | • **Preclinical** – 5/5 studies conducted in lissencephalic (rodent) species.<br>• **CAMAROS** – human participants<br>• **Interpretation** – effects of CCR5 antagonists for stroke recovery in gyrencephalic species represents a critical knowledge gap |
| (6) Tests during the awake phase of animals | No | No | • **Preclinical** – 0/5 studies reported assessing outcomes at night (i.e. awake phase of rodents)<br>• **CAMAROS** – follow-up occurs during daytime hours (i.e. awake phase of humans)<br>• **Interpretation** – future preclinical studies should conduct behavioural testing during the awake phase to better align with human testing conditions |

**Table 4.** Alignment of included preclinical studies with additional STAIR / SRRR items and CAMAROS trial parameters.

| Recommendation | Preclinical evidence | CAMAROS alignment | Notes |
| --- | --- | --- | --- |
| (1) Testing in both permanent and transient occlusion models | Yes | Yes | • **Preclinical** – 2/5 studies permanent occlusion, 2/5 transient occlusion, 1/5 hemorrhagic<br>• **CAMAROS** – participants with ischemic anterior circulation stroke, with or without reperfusion treatments eligible for enrollment |
| (2) Monitoring of treatment effects on physiological parameters, including temperature, both during and after ischemia | No | NA | • **Preclinical** – 1/5 studies (Takami 2002; TAK-779) monitored and provided data on temperature for an extended period post-stroke<br>• **CAMAROS** – participants will not be administered maraviroc in a cerebroprotective context |
| (3) Testing interaction with thrombolytics and other medications commonly administered in acute stroke | No | No | • **Preclinical** – 0/5 studies included assessed drug interactions<br>• **CAMAROS** – participants may or may not be exposed to thrombolytics or other concomitant medications that could interact with recovery-inducing effects of maraviroc |
| (4) Animal model should produce infarction similar in relative size and location to that observed in humans | Yes | Yes | • **Preclinical** – 1/5 studies (Joy 2019; maraviroc) produced infarcts in motor regions of control animals that were less than 25% of hemispheric volume<br>• **CAMAROS** - participants with ischemic anterior circulation stroke with hemiparesis requiring rehabilitation eligible for enrollment |
| (5) For studies claiming recovery effects, analysis of infarct volume should be performed to show equivalency of injury in the treated and stroke control groups | No | NA | • **Preclinical** – 0/1 experiments in late sub-acute/early chronic post-stroke phase (Joy 2019) demonstrated that each treatment group had equivalent baseline stroke volumes. This could potentially confound observed behavioral effects |
| (6) For studies claiming recovery effects, tissue repair / neuroplasticity processes should be quantified and directly related to behavioral recovery | Yes | NA | • **Preclinical** – 1/5 studies (Joy 2019) demonstrated maraviroc association with dendritic spine preservation, axonal projections to contralateral cortex, reduced inflammatory response, and upregulation of CREB / DLK signalling<br>• **CAMAROS** – outcomes to assess mechanism of recovery are not included |
| (7) For studies claiming recovery effects, initial impairment with spontaneous recovery that plateaus significantly below pre-stroke performance | No | NA | • **Preclinical** – 1/5 studies assessed long-term outcomes (Joy 2019). Large variability in initial levels of impairment across experiments. Control animals demonstrate limited spontaneous recovery or atypical worsening of performance across time in multiple experiments.<br>• **CAMAROS** – placebo participants will receive 8-week exercise program making assessment of spontaneous recovery difficult |
| (8) Behavioral effects should be assessed across a battery of domains, including both skilled and spontaneous upper limb and hindlimb use | No | No | • **Preclinical** – 2/5 studies included battery of behavioral tasks, but no study assessed effects on upper extremity skilled reaching / grasping or alteration of movement kinematics<br>• **CAMAROS** – two primary outcomes: 1) upper extremity assessment using Fugl-Meyer; 2) change in 10-meter walk test score. Preclinical outcomes have limited relevance to Fugl-Meyer |
| (9) Mechanism of action should be assessed through gain of function / loss of function studies and directly associated to behavioral effects | Yes | NA | • **Preclinical** – 1/5 studies (Joy 2019) demonstrated that knockdown of DLK signalling using small hairpin DLK is able to block beneficial behavioral effects of CCR5 knockdown, providing a possible mechanism for stroke recovery-inducing effects<br>• **CAMAROS** – outcomes to assess mechanism of recovery are not included |
| (10) Testing interactions of treatment with clinically inspired best practice, such as training or enrichment | No | No | • **Preclinical** – 0/5 studies included behavioral therapy / rehabilitative interventions<br>• **CAMAROS** – all participants (including placebo) will participate in an 8-week<br>exercise program |

For CCR5 antagonists as a post-stroke cerebroprotectant, the overall body of evidence satisfied all five STAIR XI domains assessing ‘candidate treatment qualification’ (Table 3). Overall, a range of doses and clinically relevant administration times for cerebroprotection were evaluated across a variety of motor and cognitive behavioural domains. All studies tested both behavioural and histological outcomes and demonstrated cerebroprotective effects, but most studies failed to measure and control post-stroke temperature, which could potentially confound the observed cerebroprotection (Table 4).^21^ Most histological measurements were also assessed at <72 hours, which could confound the observed cerebroprotective effects if cell death was merely delayed.^21^ For CCR5 antagonists as a post-stroke recovery-inducing treatment, one experiment assessed the effects of initiating CCR5 administration in a similar post-stroke phase as the CAMAROS trial. This experiment (Joy et al.)^6^ did not demonstrate that each treatment group had equivalent baseline stroke volumes, which may potentially confound observed behavioural effects. Furthermore, the maximum dose used in mice (100 mg/kg) was not sufficient to attain cerebrospinal fluid levels of maraviroc observed in humans using the CAMAROS dosing regime (mice: 13.8±5.4 ng/mL; humans 33.6–60 ng/mL).^6^

Areas of concern were identified in all STAIR XI domains assessing ‘preclinical assessment and validation’ (Table 3). Although adult animals were used in most of the preclinical studies, the reported weights and ages of these animals corresponded to young adults, rather than the aged adults that better represent the stroke population. All experiments used only male rodents that were free of common stroke comorbidities. It was also unclear if behavioural testing was performed during the inactive circadian phase or active (dark) phase, which could result in confounding if CCR5 antagonists affect arousal of animals during their inactive period.^22^ Furthermore, none of the studies had their protocols or results directly replicated by an independent laboratory, or across multiple sites in a multilaboratory study.^9,23^ Regarding stroke recovery, no studies assessed behavioural effects on upper extremity skilled reaching / grasping or potential interactions of CCR5 antagonists with rehabilitative therapies or established recanalization procedures (Table 4).^24–27^ These elements are highly relevant to the CAMAROS trial, as one of the primary outcomes of this trial is upper extremity performance on the Fugl-Meyer and maraviroc administration will be paired with an 8-week exercise program. These findings were supported by the PRIMED^2^ tool, which resulted in a Readiness for Translation Score of “medium” (on a scale of “low”, “medium”, “high”). This tool highlighted similar promising elements, as well as weaknesses, as our analysis above (Table 3, Table 4), identifying the limited preclinical evidence for the effects of CCR5 antagonists in clinically relevant sexes, ages, species, and disease comorbidities, without sufficient dose-response information to inform trials (Appendix 2-table 1). Overall, our assessments highlight a variety of knowledge gaps, potential confounding factors, and areas of misalignment between the preclinical evidence and clinical trial parameters that could be improved with further preclinical experimentation.

## Discussion

The overall body of preclinical evidence for CCR5 antagonists in stroke demonstrates potential acute cerebroprotection with corresponding impairment reduction, as well as improved functional recovery in the sub-acute/early chronic phase. Our systematic review also highlights evidence gaps that could impact successful clinical translation of CCR5 antagonists. Our analysis of 10 independent experiments, identified that acute administration of CCR5 antagonists within the first 24 hours post-stroke was associated with a marked reduction in infarct volume. This cerebroprotective reduction of infarct volume did not significantly vary based on treatment dose or any other experimental characteristics.^3,4,6^ Overall, the majority of behavioural effects appeared to be in a positive direction, but the low number of included studies precluded meta-analysis of these results. Indeed, no individual behavioural experiment included more than ten CCR5-treated animals, and given the wide range of dosages, timings, routes, stroke models, and rodent strains involved, the certainty of these findings is limited and should be interpreted cautiously. Pooling data across heterogenous experimental designs, animal/stroke models, and treatment parameters, as we have done with the infarct volume analysis in the present study, can introduce variability that increases the risk of overestimating or underestimating the true effect of the intervention.^28^ Treatment effects observed across model systems and therapeutic compounds may represent different biological mechanisms. Despite this potential limitation, meta-analysis can provide valuable insights, especially in preclinical settings where the sample sizes of individual studies may be too small to detect significant effects on their own. In these cases, pooling data across studies can help identify overarching estimates of benefits and harm, highlight subgroups of interest, and help guide areas of future research. As described in the results above, we attempted to mitigate the risks of inappropriate data pooling through careful investigation of heterogeneity, subgroup analyses, and differentiation between outcomes where we felt that meta-analytic pooling was (infarct volume) and was not (behavioural outcomes) appropriate. Overall, we believe that our results indicate that further investigation is warranted to determine the optimal timing of administration and behavioural domains under which CCR5 antagonists exhibit the strongest post-stroke cerebroprotective and recovery-inducing effects.

Despite the positive direction of treatment effects across all studies of CCR5 antagonists, we found a substantial risk of bias in the underlying studies.^15^ As is commonly observed in the preclinical literature, all studies either did not adequately report their randomization / blinding methods and exhibited evidence of selective / incomplete reporting.^16–18^ Such features are associated with biased overestimations of preclinical treatment efficacy, which raises further concerns about the reliability and validity of the present findings.^29–31^ Future studies should carefully incorporate all elements of the ARRIVE 2.0 guidelines to help ensure that results are transparently reported and improve confidence in the findings.^32^

Comprehensiveness of the preclinical evidence for CCR5 antagonists was assessed in relation to STAIR and SRRR consensus recommendations.^9,11,19,20^ These recommendations aim to provide investigators and regulators with “assurance that the candidate treatment shows signals of efficacy and safety, before embarking on an expensive clinical development program”.^11^ The included studies provide good initial evidence for acute cerebroprotection, as well as mechanistic and behavioural evidence for enhanced recovery in the late sub-acute/early chronic post-stroke phase. However, demonstration of efficacy under a wider range of conditions, such as in aged animals, females, animals with stroke-related comorbidities, more clinically relevant timing of dose administrations, or in conjunction with rehabilitative therapies are necessary to provide further confidence in these findings. In addition, all studies used unique doses, timings, and outcomes, so independent replication of the most promising study parameters would further increase certainty in the evidence.^9,11^ Future preclinical research should aim to address these evidence gaps to further increase the clinical relevance and comprehensiveness of evidence for CCR5 antagonists in stroke.

In relation to the ongoing CAMAROS trial assessing maraviroc in the subacute post-stroke phase,^7^ the most relevant preclinical evidence comes from one experiment within the Joy et al. study^6^ where maraviroc was initially administered at three to four weeks post-stroke and continued daily for 11 weeks. This experiment demonstrated that administration of maraviroc in the late sub-acute/early chronic post-stroke phase improved functional recovery on the grid walk, but not cylinder, task. These experimental conditions could potentially align with the putative therapeutic window and outcomes of interest being assessed in CAMAROS (e.g. 10-minute walk test co-primary outcome).^7^ However, caution is warranted as this pivotal supporting preclinical evidence is based on a low sample size (n=9). Moreover, potential differences in dosing, severity of infarct, concomitant rehabilitative therapy, and other factors discussed above could influence the degree to which these results successfully translate to the clinical environment. Finally, clinically relevant secondary outcomes in the CAMAROS trial, such as cognitive and emotional domains as measured by the Montreal Cognitive Assessment (MoCA) and Stroke Aphasia Depression Questionnaire (SADQ) were not modelled in the preclinical literature. Although one study included a cognitive outcome, the other treatment parameters of this study were not aligned to the CAMAROS trial.^5^ Future preclinical studies should assess a more diverse and comprehensive battery of clinically relevant behavioural tasks, which could be based on the range of outcomes employed in the CAMAROS trial, or those found in the SRRR recommendations.^9^ Nevertheless, the Joy et al., study provides a plausible biological mechanism and “proof of concept” for how CCR5 antagonism might enhance neuroplasticity that improves functional recovery after stroke.

Our present synthesis of the preclinical evidence for CCR5 antagonists used novel approaches to increase its utility for assessing certainty of the findings and identification of knowledge gaps. First, we engaged patients with lived experiences of stroke throughout the review process to ensure that our research questions, outcomes, and interpretations aligned with the priorities of the ultimate end-user of stroke research. Second, we incorporated consensus recommendations for both preclinical cerebroprotection and recovery research as an evidence evaluation tool, which we found often aligned with the priorities of our patient panel.^9,11,19,20^ This guided assessment of the alignment of preclinical evidence with parameters of ongoing clinical trials, as well as appraisal of comprehensiveness of the preclinical evidence with a focus on translational validity^33^ rather than only internal validity and risk of bias.^15^ We also used this method to provide concrete avenues for future preclinical studies to close knowledge gaps and improve certainty in the effects of CCR5 antagonists under clinically relevant experimental conditions. Similar approaches should be considered by future preclinical systematic reviews to improve interpretation of the preclinical evidence from a translational perspective.

In conclusion, CCR5 antagonists show promise in preclinical studies for stroke cerebroprotection, corresponding reduction in impairment, as well as improved functional recovery related to neural repair in the late sub-acute/early chronic phase. However, high risk of bias and the limited (or no) evidence in clinically relevant domains underscore the need for more rigorous and transparent preclinical research to further strengthen the current preliminary evidence available in the literature. Addressing these concerns will not only enhance the reliability of preclinical evidence but also better inform the design and execution of clinical trials of CCR5 antagonists, such as the ongoing CAMAROS trial.^7^ The integration of expert recommendations, such as STAIR and SRRR, should guide future preclinical investigations and synthesis of the body of preclinical evidence in stroke recovery research.^9,11^ Our present approach serves as a template by which the preclinical evidence supporting translation to clinical trials can be weighed when justifying early clinical trials of novel interventions for stroke cerebroprotection and recovery.

## Materials and methods

We registered the review protocol on the International Prospective Register of Systematic Reviews (CRD42023393438).^34^ The findings are reported in accordance with the Preferred Reporting Items for Systematic Reviews and Meta-Analyses (see attached Reporting Standards Document)^35^ and Guidance for Reporting the Involvement of Patients and the Public (Appendix 3-table 1).^36^

### Engagement with patient panel - individuals with lived experience of stroke

A panel of eight patients and caregivers with lived experience of a stroke informed project development (e.g., research question development, review protocols, search strategy development) and were actively involved in the research conduct (screening, data extraction, analysis, interpretation). Monthly meetings occurred with the patients and caregivers to provide educational sessions of background knowledge of preclinical stroke, systematic review conduct, and discuss research findings as the review progressed. We co-developed a terms of reference document *a priori* to document details of the engagement (i.e., roles, responsibilities, expectations, project goals, etc.). The patient partners co-developed the research question to address patient interests, including chronic stroke recovery (i.e., the panel was particularly interested in evaluating the effects of extended drug administration), consideration of physical therapy in tandem with drug administration, inclusion of stroke-relevant comorbidities, spasticity, and motor and cognitive outcomes. Co-authorship and financial compensation were agreed upon with the patients and caregivers and offered as a method of acknowledgement according to the Canadian Institutes of Health Research (CIHR) Strategies for Patient Oriented Research (SPOR) Evidence Alliance Patient Partner Appreciation Policy.

### Eligibility criteria

- *Animals:* Any preclinical *in vivo* animal models of adult stroke were included. Human, invertebrate, *in vitro*, *ex vivo*, and neonatal animal studies were excluded.
- *Model:* Focal ischaemic or intracerebral haemorrhagic stroke models were included, while animal models of global ischaemia were excluded.
- *Intervention:* Studies administering a CCR5 antagonist drug (e.g., maraviroc, D-Ala-Peptide T-Amide (DAPTA), Takeda 779 (TAK-779)). Study arms in which CCR5 was genetically manipulated (e.g., CCR5 knockout strain) were excluded.
- *Comparator:* Vehicle-treated control groups where stroke was induced. CCR5 antagonist control groups without stroke were excluded.
- *Outcome:* Studies reporting at least one of the following: infarct size, behavioural tests, mortality, adverse events, and spasticity were included.
- *Study design and publication characteristics:* Controlled interventional studies (randomized, pseudo-randomized, or non-randomized) published as full journal articles in any language or year were included. Abstracts, review articles, opinion-based letters/editorials, and unpublished grey literature were excluded.

### Information sources and search strategy

An information specialist with experience in systematic searches of the preclinical literature developed a comprehensive search strategy based on a previously published strategy for identifying animal experimentation studies (Appendix 4).^37^ The search strategy underwent peer-review using the Peer Review of Electronic Search Strategies (PRESS) checklist.^38^ We searched MEDLINE (OVID interface, including In-Process and Epub Ahead of Print), Web of Science, and Embase (OVID interface). The search was conducted October 25, 2022, to align with the listed launch date of the CAMAROS trial (September 15, 2022). Our intention in doing so was to collate and assess all preclinical evidence that could have feasibly informed the clinical trial. We sought to assess the comprehensiveness of evidence and readiness for translation of CCR5 antagonist drugs at the time of their actual translation into human clinical trials, as well as the alignment of the CAMAROS trial design to the existing preclinical evidence base.

### Selection process and data collection

We deduplicated citations and uploaded them into DistillerSR® (Evidence Partners, Ottawa, Canada). Two reviewers independently screened citations by title and abstract using an accelerated method (one reviewer required to include, two reviewers required to exclude). We then screened and extracted data from full-text articles in duplicate. Graphical data was extracted using Engauge Digitizer.^39^ A third reviewer with content expertise in preclinical stroke studies audited all data extraction. Conflicts between reviewers were resolved by consensus discussion. See Appendix 5 for the complete list of data extraction elements.

### Effect measures and data synthesis

We performed quantitative analyses using the R (version 4.1.2) “metafor” package (version 4.0.0)^40^ with inverse variance random effects modelling. We expressed continuous outcome measures as standardized mean differences (SMDs) with 95% confidence intervals (CIs) and assessed statistical heterogeneity of effect sizes using the Cochrane I² statistic.^41^ This was necessary due to the variety of outcome measures and measurement scales used across studies, which is a common feature of preclinical systematic reviews. Sensitivity analyses were performed using original measurement scales where possible. From patient partner input, subgroups were analyzed based on timing / dose / route of intervention, stroke model, stroke type, species, type of behavioural outcome (i.e., motor, cognitive), and comorbidities.^42^ Our planned subgroup analyses on study quality, specific regions/areas of the brain, and post-stroke rehabilitation paradigms were not performed due to insufficient number of studies. We did not assess publication bias using Egger plots due to fewer than 10 studies being included in the analysis, as per Cochrane recommendations.^43^

### Risk of bias assessment

Two independent reviewers used the Systematic Review Centre for Laboratory Animal Experimentation (SYRCLE) risk of bias tool to assess each study as having a “Low Risk”, “Unclear Risk”, “Some Concerns of Risk”, or “High Risk” across domains such as randomization, blinding and outcome reporting.^15^ “Some Concerns of Risk” indicated reporting of a domain (i.e., randomization), but lacking methodological details. This differed from “Unclear Risk” where there was no mention of the domain in the study.

### Comprehensiveness of preclinical evidence and alignment with clinical trials

We assessed comprehensiveness of the *overall body* of preclinical evidence for CCR5 antagonists as a cerebroprotective treatment using The Stroke Treatment Academic Industry Roundtable (STAIR) I, VI, and XI Consolidated Recommendations.^11,19,20^ These recommendations encompass “candidate treatment qualification” (e.g., dose, timing of dose, outcomes) and “preclinical assessment and validation” (e.g., age, sex, sample size, animal type).^11^ We excluded domains that were redundant with risk of bias (e.g., randomization, blinding, etc.) and included additional items relevant to stroke recovery studies from the Stroke Recovery and Rehabilitation Roundtable (SRRR) Translational Working Group consensus recommendations.^9^ Two reviewers extracted data to determine if the overall evidence across studies satisfied each of the STAIR and SRRR recommendations. A third reviewer audited this analysis. We then assessed alignment of existing preclinical evidence with an ongoing clinical trial of CCR5 antagonists for stroke (CAMAROS, NCT04789616).^7^

### Protocol deviations

Additionally, a list of outcomes used to determine CCR5’s potential mechanisms of action were extracted (Appendix 1-figure 1, Appendix 1-table 1) based on feedback from individuals that reviewed the initial manuscript drafts. We also incorporated an additional assessment using the PRIMED^2^ tool for assessing the readiness of stroke cerebroprotection therapies to be translated to clinical trials based on the body of preclinical evidence (Appendix 2-table 1).^44^

## Acknowledgements

The authors thank Hannah Laquerre for her assistance with figure generation. Appendix 1-figure 1 was created with BioRender.com.

## Sources of Funding

This study was funded by a Social Accountability Grant from the University of Ottawa’s Faculty of Medicine. MSJ was supported by the Canadian Institutes of Health Research (CIHR) and Vanier Canada Graduate Scholarship (CGV-186957). The funders were not involved in the study design, collection, analysis and interpretation of data, writing the manuscript, or in the decision to submit the article for publication. MML is supported by The Ottawa Hospital Anesthesia Alternate Funds Association, a University of Ottawa Junior Research Chair, and the Canadian Anesthesiologists’ Society Career Scientist Award.

## Disclosures

None.

## Data Availability

All data generated or analysed during this study are included in the manuscript and supporting files. Source data and code to generate Figures 2 and 3 have been provided at the following DOI: https://doi.org/10.17605/OSF.IO/5KGT6

## Non-standard Abbreviations and Acronyms

CAMAROS: Canadian Maraviroc RCT to Augment Rehabilitation Outcomes After Stroke
CCR5: C-C chemokine receptor type 5
DAPTA: D-Ala-Peptide T-Amide
SRRR: Stroke Recovery and Rehabilitation Roundtable
STAIR: Stroke Treatment Academic Industry Roundtable
SYRCLE: Systematic Review Centre for Laboratory Animal Experimentation
TAK-779: Takeda 779

## Appendix 1. Potential mechanisms of action

**Appendix 1-figure 1.**
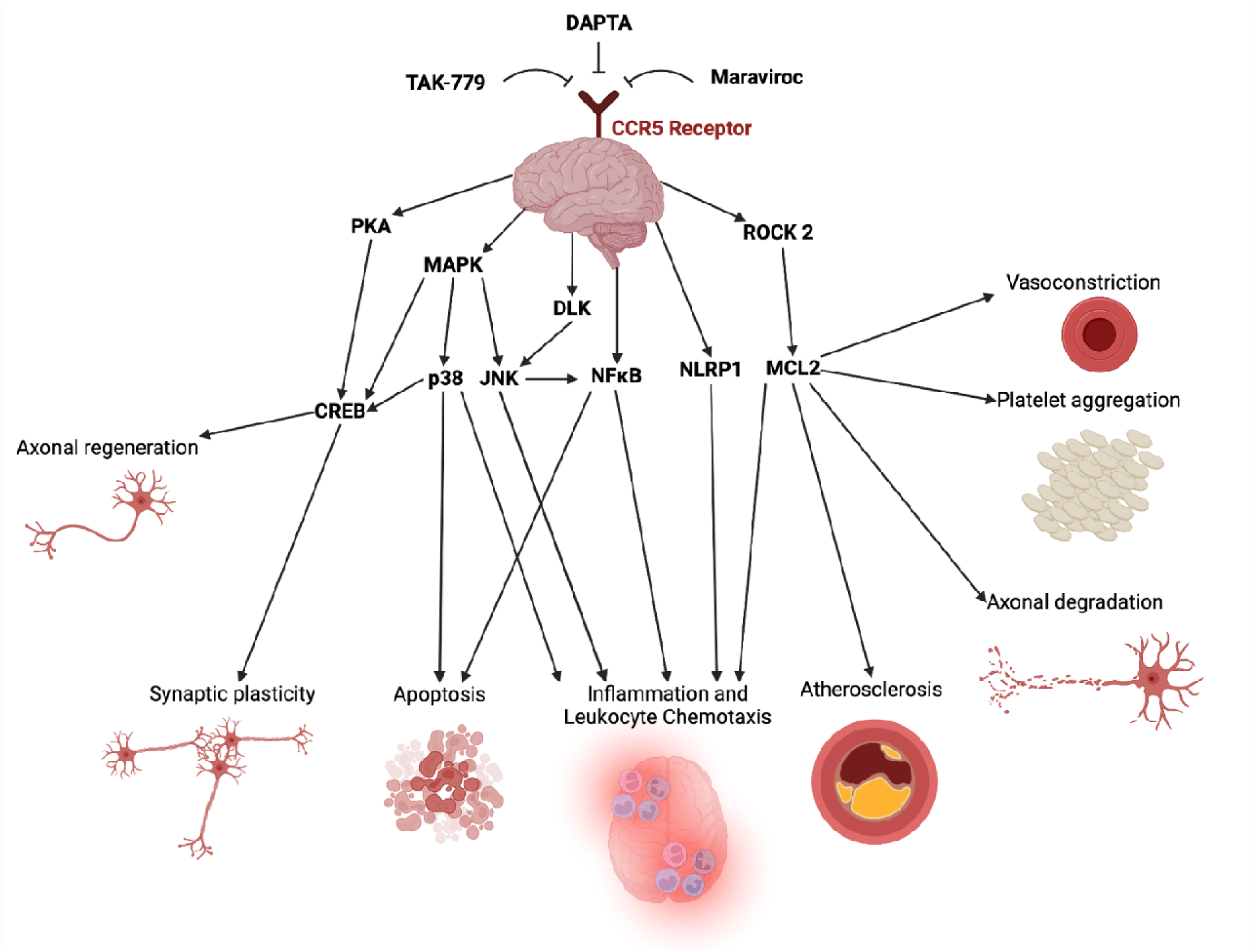
Potential mechanistic pathways and proposed domains of biological activity underlying CCR5 antagonist’s cerebroprotective and neural repair effects post-stroke described in the included studies. A list of mechanistic evidence supporting these pathways was extracted from included studies in Appendix 1-table 1. Created with BioRender.com.

**Appendix 1-table 1.** List of outcomes used to determine potential mechanisms of action, by study.

| Paper | Mechanistic evidence presented |
| --- | --- |
| Takami 2002 –<br>*TAK-779* | - Infarct volume reduction (reduced by TAK-779)<br>- MPO + Mac-1-alpha antibody used to stain infiltrating neutrophils/macrophages/microglia (Mac-1-alpha reduced by TAK-779 / MPO not decreased) – neutrophils the same, macrophages/microglia reduced |
| Li 2016 –<br>*DAPTA* | - Cell death reduction (TUNEL / H&E)<br>- Western blot – ROCK2 / P-MLC2 reduced (by DAPTA) |
| Joy 2019 –<br>*maraviroc* | - CCR5 knockdown FACS isolated neuronal assay -> CREB / pCREB / DLK significantly increased. Erk(p44), GAP43, SAP/JNK1+2 increased, but not significantly. P38 decreased, but not significantly<br>- 2-photon imaging - CCR5 knockdown + maraviroc -> increased dendritic spine count, fewer lost<br>- BDA axonal quantification – CCR5 knockdown + maraviroc -> increased bihemispheric sprouting<br>- GFAP / IBA-1 immunostaining – CCR5 knockdown + maraviroc -> reduced immunoreactivity. Decreased Ly6C (neutrophil) and Ly6Clow (macrophage) recruitment |
| Yan 2021 –<br>*maraviroc* | - Cell death reduction (TUNEL / FJC / cleaved caspase 1 / Nissl)<br>- Western blot – maraviroc -> increased PK A-Calpha / pCreb -> decreased NLRp1 / cleaved caspase 1 / IL-1B / apoptosis-associated speck protein / N-gasdermin D / IL-18<br>- Western blot -> effects reversed by 666-15<br>- Western blot -> rCCL5 shows reversed effects from maraviroc -> rCCL5 + cAMP reverses those effects |
| Chen 2022 –<br>*maraviroc* | - Infarct volume / Cell death reduction (TTC / TUNEL)<br>- Western blot – maraviroc -> increased Bcl2:BAX ratio -> increased IκBα, -> decreased P-IκBα / P-P65 / P-P38 / P-JNK<br>- Western blot – P-JNK decreased, Anisomycin reverses the effect on P-JNK<br>- qRT-PCR, ELISA -> reduced IL-1B / IL-6 / TNF-alpha |

**Appendix 2-table 1.**
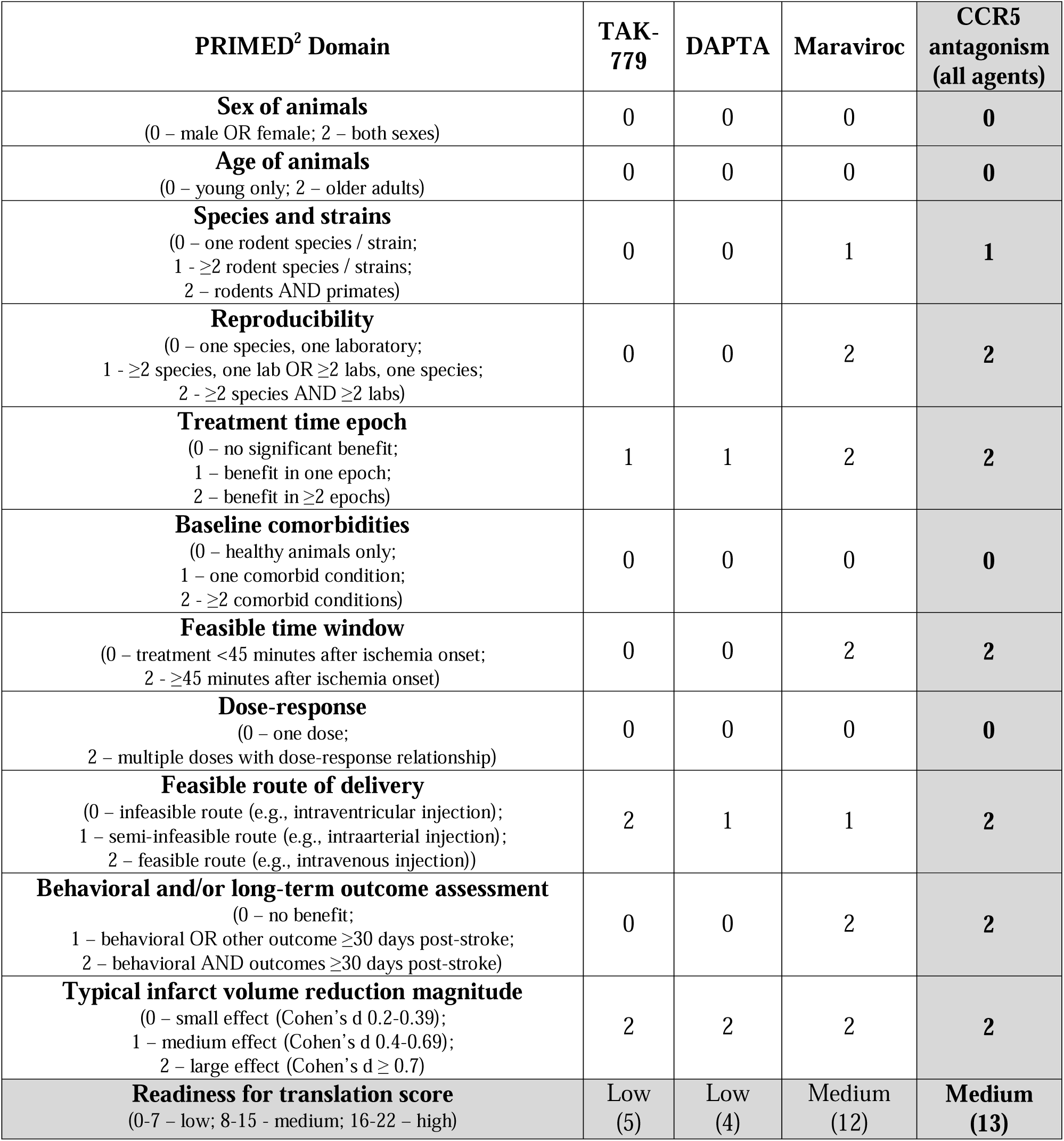
Readiness of CCR5 antagonists for translation based on the PRIMED2 tool.

| **PRIMED<sup>2</sup> Domain** | **TAK-779** | **DAPTA** | **Maraviroc** | **CCR5 antagonism (all agents)** |
| --- | --- | --- | --- | --- |
| **Sex of animals**<br>(0 – male OR female; 2 – both sexes) | 0 | 0 | 0 | **0** |
| **Age of animals**<br>(0 – young only; 2 – older adults) | 0 | 0 | 0 | **0** |
| **Species and strains**<br>(0 – one rodent species / strain;<br>1 – $\geq 2$ rodent species / strains;<br>2 – rodents AND primates) | 0 | 0 | 1 | **1** |
| **Reproducibility**<br>(0 – one species, one laboratory;<br>1 – $\geq 2$ species, one lab OR $\geq 2$ labs, one species;<br>2 – $\geq 2$ species AND $\geq 2$ labs) | 0 | 0 | 2 | **2** |
| **Treatment time epoch**<br>(0 – no significant benefit;<br>1 – benefit in one epoch;<br>2 – benefit in $\geq 2$ epochs) | 1 | 1 | 2 | **2** |
| **Baseline comorbidities**<br>(0 – healthy animals only;<br>1 – one comorbid condition;<br>2 – $\geq 2$ comorbid conditions) | 0 | 0 | 0 | **0** |
| **Feasible time window**<br>(0 – treatment <45 minutes after ischemia onset;<br>2 – $\geq 45$ minutes after ischemia onset) | 0 | 0 | 2 | **2** |
| **Dose-response**<br>(0 – one dose;<br>2 – multiple doses with dose-response relationship) | 0 | 0 | 0 | **0** |
| **Feasible route of delivery**<br>(0 – infeasible route (e.g., intraventricular injection);<br>1 – semi-infeasible route (e.g., intraarterial injection);<br>2 – feasible route (e.g., intravenous injection)) | 2 | 1 | 1 | **2** |
| **Behavioral and/or long-term outcome assessment**<br>(0 – no benefit;<br>1 – behavioral OR other outcome $\geq 30$ days post-stroke;<br>2 – behavioral AND outcomes $\geq 30$ days post-stroke) | 0 | 0 | 2 | **2** |
| **Typical infarct volume reduction magnitude**<br>(0 – small effect (Cohen's d 0.2-0.39);<br>1 – medium effect (Cohen's d 0.4-0.69);<br>2 – large effect (Cohen's d $\geq 0.7$) | 2 | 2 | 2 | **2** |
| **Readiness for translation score**<br>(0-7 – low; 8-15 – medium; 16-22 – high) | Low<br>(5) | Low<br>(4) | Medium<br>(12) | **Medium**<br>**(13)** |

**Appendix 3-table 1.** GRIPP2 Short Form Checklist.

| Section and topic | Item |
| --- | --- |
| **1: Aim** | To conduct a preclinical systematic review assessing the effects of C-C chemokine receptor type 5 (CCR5) antagonists on motor and cognitive impairment following stroke. To collaborate with a panel of patients and caregivers with lived experience of stroke throughout the development and conduct of the preclinical systematic review. |
| **2: Methods** | A panel of eight patients and caregivers with lived experience of stroke was recruited to join the research team through the Heart & Stroke Foundation and the Patient and Family Advocacy Program at The Ottawa Hospital. Recruitment advertisements were distributed to both organizations. The patients and caregivers were involved in developing the research question and the protocol (i.e. elements of the AMICO question presented in the Methods section); identifying data items for extraction; conduct of the review including screening and extraction; interpreting the review results; and contributed to edits to the final manuscript. Patients and caregivers attended monthly virtual meetings. These meetings provided background knowledge of preclinical stroke, systematic review conduct, and discussed research findings as the review progressed. Patients provided insights into their lived experiences with stroke and helped identify priority areas that they were interested in. Patients and caregivers were offered financial compensation and co-authorship in recognition of their contributions to the research project. |
| **3: Results** | Patient engagement contributed to the study in several ways, including:<br>- Informing the research question with the patient partner experience: patient partners have lived experience of stroke and provided patient priorities to analyze in our review<br>- Refining our primary and secondary outcomes to include patient priorities<br>- Developing of the protocol to include non-technical language<br>- Editing screening and extraction forms<br>- Participating as reviewers in abstract and full-text level screening<br>- Participating as reviewers in extraction<br>- Attending an international stroke conference and participating as co-authors of an abstract presented there<br>- Interpreting analyzed data, including identifying patient important barriers and limitations to incorporate in the discussion |
| **4: Discussion** | Overall, patient engagement was successful in informing review development and conduct. Additionally, the researchers on the team learned from the patient panel. The patient panel brought a unique perspective to the planning and conduct of a preclinical systematic review. Additionally, members of the panel stated that the experience was useful for them as they gained new insights into preclinical research. |
| **5: Reflections** | Engagement was embedded within the research project from the inception to dissemination, where patient partners were members of the larger team. Patient partners directed us to outcomes for the systematic review that critical clinically to patients in the chronic phase of stroke recovery. Their participation in the conduct of the review facilitated meaningful collaborations and discussion. |

## Appendix 4. Full search strategies for MEDLINE / Embase and Web of Science

Medline: 58

Embase: 118

Web of Science: 87

Total: 230

Total after duplicates: 166 Duplicates: 64

******************************************************************************

Embase Classic+Embase <1947 to 2022 October 24>

Ovid MEDLINE(R) ALL <1946 to October 24, 2022>

1 exp animal experimentation/ or exp models, animal/ or animals/ or exp animal population groups/ or chordata/ or vertebrates/ or exp amphibians/ or exp birds/ or exp fishes/ or exp reptiles/ or mammals/ or primates/ or eutheria/ or exp artiodactyla/ or exp carnivore/ or exp cephalopoda/ or exp cetacea/ or exp chiroptera/ or exp elephants/ or exp hyraxes/ or exp insectivora/ or exp lagomorpha/ or marsupialia/ or exp monotremata/ or exp perissodactyla/ or proboscidea mammal/ or exp rodentia/ or exp scandentia/ or exp sirenia/ or exp cingulata/ or haplorhini/ or exp strepsirhini/ or exp platyrrhini/ or exp tarsii/ or catarrhini/ or exp cercopithecidae/ or exp hylobatidae/ or hominidae/ or exp gorilla gorilla/ or exp pan paniscus/ or exp pan troglodytes/ or exp pongo/ 38941083

2 (rat or rats or animal or animals or mice or in vivo or mouse or rabbit or rabbits or murine or pig or pigs or dog or dogs or bovine or fish or vertebrate or vertebrates or cat or cats or rodent or rodents or mammal or mammals or chicken or chickens or monkey or monkeys or sheep or canine or canines or porcine or cattle or bird or birds or hamster or hamsters or primate or primates or cow or cows or chick or horse or horses or avian or avians or calf or swine or swines or xenopus or turkeys or bear or bears or frog or frogs or zebrafish or goat or goats or equine or calves or poultry or macaque or macaques or mole or moles or ovine or lamb or lambs or fishes or diptera or amphibian or amphibians or snake or snakes or ruminant or ruminants or hen or hens or piglet or piglets or feline or felines or simian or simians or laevis or trout or trouts or teleost or teleosts or salmon or salmons or seal or seals or bull or bulls or ewe or ewes or hedgehog or hedgehogs or macaca or macacas or proteus or pigeon or pigeons or bat or bats or duck or ducks or chimpanzee or chimpanzees or baboon or baboons or deer or rana or ranas or carp or carps or heifer or swallow or swallows or lizard or lizards or canis or sow or sows or cynomolgus or quail or quails or reptile or reptiles or turtle or turtles or buffalo or gerbil or gerbils or boar or boars or squirrel or squirrels or oncorhynchus or mus or toad or toads or fowl or fowls or rerio or danio or ara or aras or musculus or tadpole or tadpoles or mulatta or salmo or ram or eagle or eagles or ferret or ferrets or goldfish or catfish or whale or whales or fox or foxes or ape or apes or elephant or elephants or bos or marmoset or marmosets or cod or cods or shark or sharks or wolf or eel or eels or auratus or rattus or zebra or zebras or tilapia or tilapias or gilt or camel or camels or squid or gallus or marsupial or marsupials or vole or voles or fascicularis or ovis or salmonid or salmonids or tiger or tigers or dolphin or dolphins or robin or robins or carpio or opossum or opossums or cyprinus or salamander or salamanders or felis or mink or minks or swan or swans or norvegicus or bufo or torpedo or bass or lamprey or lampreys or sus or python or pythons or tetrapod or tetrapods or shrew or shrews or lion or lions or hog or hogs or songbird or songbirds or oreochromis or starling or starlings or caprine or carassius or owl or owls or newt or newts or papio or scrofa or hare or hares or gorilla or gorillas or flounder or flounders or goose or herring or herrings or therian or buffaloes or canary or sparrow or sparrows or microtus or octopus or troglodytes or tuna or amphibia or chinchilla or chinchillas or ide or oryzias or cervus or kangaroo or kangaroos or armadillo or armadillos or callithrix or pan troglodytes or saimiri or cichlid or cichlids or donkey or donkeys or bream or char or chars or finch or raccoon or raccoons or bothrops or anguilla or perch or cricetus or seabird or seabirds or buck or bucks or naja or coturnix or salmonids or geese or minnow or minnows or raptor or raptors or merione or meriones or rodentia or elaphus or amniote or amniotes or elasmobranch or emu or emus or peromyscus or hominid or hominids or bubalus or crotalus or gull or gulls or anas or anura or lemur or lemurs or crow or crows or camelus or gibbon or gibbons or waterfowl or parrot or parrots or eels or cob or stickleback or sticklebacks or columba or mesocricetus or ambystoma or raven or ravens or gadus or penguin or penguins or orangutan or orangutans or sturgeon or sturgeons or cuniculus or aves or virginianus or cephalopod or cephalopods or cebus or sparus or tortoise or tortoises or guttata or morhua or unguiculatus or dogfish or vulpes or mallard or mallards or apodemus or alligator or alligators or oryctolagus or llama or llamas or reindeer or mustela or duckling or ducklings or wolves or sander or amazona or zebu or badger or badgers or dove or doves or ictalurus or capra or capras or equus or camelid or camelids or poecilia or mule or mules or perciformes or salvelinus or labrax or cyprinidae or ariidae or crocodile or crocodiles or fundulus or dicentrarchus or clarias or cercopithecus or chiroptera or alpaca or alpacas or pike or pikes or paralichthys or puma or pumas or didelphis or pisces or macropus or triturus or bison or bisons or epinephelus or gasterosteus or panthera or acipenser or mackerel or mackerels or tamarin or tamarins or ostrich or anolis or vervet or vervets or wallaby or glareolus or beaver or beavers or dromedary or catus or killifish or pimephales or promelas or aotus or phoca or panda or pandas or porpoise or porpoises or myotis or yak or yaks or agkistrodon or vipera or otter or otters or turbot or turbots or squamate or carnivora or mullet or mullets or hawk or hawks or taeniopygia or seahorse or seahorses or poecilia reticulata or falcon or falcons or prosimian or prosimians or parus or perca or fingerling or fingerlings or antelope or antelopes or tupaia or passeriformes or sepia or saguinus or coyote or coyotes or pongo or meleagris or reptilia or lepus or psittacine or hagfish or warbler or warblers or russell’s viper or russell’s vipers or smolt or smolts or budgerigar or sardine or sardines or cavia or cavias or hyla or pleurodeles or siluriformes or great tit or great tits or guppy or bonobo or bonobos or rutilus or trichosurus or muridae or phodopus or channa or squalus or lynx or sturnus or petromyzon or vitulina or monodelphis or cuttlefish or adder or adders or lepomis or canaria or gambusia or guppies or xiphophorus or flatfish or koala or koalas or labeo or stingray or stingrays or chelonia or lampetra or spermophilus or crocodilian or passer domesticus or sciurus or artiodactyla or ranidae or corvus or necturus or platypus or canaries or bovid or lagopus or trimeresurus or gariepinus or marten or martens or drosophilidae or mugil or sunfish or porcellus or cypriniformes or alouatta or scophthalmus or anser or electrophorus or putorius or iguana or iguanas or lama or lamas or takifugu or circus or eptesicus or flycatcher or galago or galagos or trachemys or lungfish or characiformes or shorebird or shorebirds or giraffe or giraffes or micropterus or scyliorhinus or cichlidae or loligo or porcupine or porcupines or chub or chubs or solea or pleuronectes or hylidae or viperidae or echis or sorex or anchovy or lagomorph or ostriches or vulture or vultures or whitefish or araneus or jird or jirds or tern or esox or drake or drakes or elapidae or gallopavo or chordata or myodes or caretta or serinus or grouse or misgurnus or meles or blackbird or blackbirds or coregonus or bobwhite or bobwhites or heteropneustes or mammoth or mammoths or turdus or rhinella or ateles or characidae or clupea or bungarus or brill or struthio camelus or sloth or sloths or pteropus or sculpin or anthropoids or pollock or pollocks or morone or pan paniscus or litoria or chipmunk or chipmunks or balaenoptera or marmota or melopsittacus or hyrax or lemming or lemmings or halibut or hylobates or lates or caiman or caimans or sigmodon or stenella or barbel or barbels or sterna or parakeet or parakeets or phocoena or leptodactylus or canidae or buteo or harengus or gopher or gophers or marmot or marmots or gosling or goslings or platichthys or gar or gars or sebastes or marsupialia or notophthalmus or gazelle or gazelles or insectivora or paridae or felidae or russula or galliformes or bombina or colobus or echidna or echidnas or seabass or syncerus or plaice or blue tit or blue tits or pagrus or catfishes or cetacea or barbus or cygnus or ficedula or chamois or colubridae or perches or coelacanth or fitch or urodela or cynops or martes or halichoerus or aix or salmonidae or leuciscus or magpie or magpies or silurus or whiting or whitings or anseriformes or colinus or rhea or chlorocebus or octodon or acinonyx or mouflon or mouflons or ibex or tetraodon or bufonidae or equidae or jackal or cephalopoda or dendroaspis or glama or muskrat or muskrats or sable or sables or wildebeest or streptopelia or albifrons or vespertilionidae or woodpecker or woodpeckers or muntjac or muntjacs or archosaur or branta or cricetulus or megalobrama or poeciliidae or desmodus or snakehead or snakeheads or tench or teal or teals or bandicoot or bandicoots or apteronotus or phyllostomidae or crocidura or buzzard or buzzards or larimichthys or cercocebus or pipistrellus or erithacus or impala or impalas or rousettus or haddock or haddocks or tinca or ratite or calidris or cynoglossus or hypophthalmichthys or bullock or bullocks or dromedaries or alectoris or filly or salamandra or cingulata or bitis or grus or ammodytes or macaw or macaws or hypoleuca or sapajus or cyprinodontiformes or hippopotamus or pelophylax or capybara or capybaras or weasel or weasels or cairina or cynomys or lutra or cockatoo or cockatoos or lachesis or lagomorpha or rupicapra or daboia or orang utan or orang utans or platyrrhini or charadriiformes or micrurus or psittaciformes or spalax or loris or mustelidae or sylvilagus or vitticeps or cockatiel or mustelus or cottus or erythrocebus or dipodomys or platessa or callicebus or loricariidae or catostomus or cuneata or cyanistes or cyprinodon or sigmodontinae or elasmobranchii or trichechus or sauropsid or xenarthra or dormouse or perissodactyla or nautilus or cirrhinus or gulo or tragelaphus or merula or numida or sciaenidae or cerastes or sciuridae or gibbosus or octopuses or eland or elands or phyllomedusa or pogona or walrus or agamidae or leptodactylidae or ridibundus or leontopithecus or anteater or anteaters or pelodiscus or cebidae or columbianus or pelteobagrus fulvidraco or hominoidea or mandrillus or zonotrichia leucophrys or agama or gobiocypris or bearded dragon or bearded dragons or sarotherodon or talpa or discoglossus or hagfishes or sphenodon or gudgeon or amphiuma or aythya or tenrec or tenrec or hominidae or risoria or salamandridae or camelidae or columbiformes or latimeria or plover or plovers or afrotheria or falco sparverius or polecat or polecats or crotalinae or salvadora or tarsier or lucioperca or anchovies or lungfishes or terrapin or dromaius novaehollandiae or lateolabrax or eigenmannia or pelamis or theropithecus or murinae or gander or gymnotus or pseudacris or gymnophiona or gymnotiformes or laticauda or falconiformes or dugong or dugongs or pintail or pintails or rook or rooks or lasiurus or catshark or catsharks or micropogonias or red junglefowl or paddlefish or ophiophagus or hollandicus or nymphicus or pimelodidae or aepyceros or cobitidae or strigiformes or cobitis or dormice or alytes or calloselasma or guanaco or phasianidae or round goby or trichogaster or catarrhini or eelpout or eelpouts or galaxias or gaur or pungitius or suslik or susliks or flatfishes or percidae or caprinae or todarodes or osmerus or ameiurus or anthropoidea or castor canadensis or pouting or poutings or tetraodontiformes or arvicolinae or siamang or siamangs or castor fiber or nomascus or red knot or red knots or syngnathidae or iguanidae or eretmochelys or ursidae or callimico or columbidae or microhylidae or anaxyrus or menidia or pipistrelle or greylag or pipidae or scandentia or bowfin or bowfins or dendrobatidae or zenaida or bushbaby or harrier or harriers or macropodidae or pygerythrus or clupeidae or odorrana or corvidae or jerboa or jerboas or canutus or hylobatidae or clupeiformes or great cormorant or great cormorants or scorpaeniformes or chondrostean or garfish or proboscidea or psetta or diapsid or serotinus or tetrao or walruses or carcharhiniformes or leucoraja or pumpkinseed or dosidicus or acipenseriformes or daubentonii or emberizidae or gadiformes or hyraxes or stizostedion or wolverine or wolverines or lissotriton or acanthurus or centrarchidae or gloydius or laurasiatheria or limosa or psittacula or leporidae or proteidae or zander or zanders or arapaima or bagridae or cyprinodontidae or mithun or pandion or jackdaw or jackdaws or procyonidae or carus or jaculus or salmoniformes or common sole or common soles or protobothrops or calamita or brachyteles or trionyx or turdidae or boidae or luscinia or pugnax or euarchontoglires or saithe or saithes or symphalangus or aardvark or aardvarks or oystercatcher or oystercatchers or arius or corydoras or poacher or poachers or aurochs or cebuella or crecca or lemuridae or sirenia or lemmus or perdix or glires or lepidosaur or muskox or deinagkistrodon or pholidota or holocephali or cercopithecinae or clariidae or agapornis or doryteuthis or tyrannidae or dicroglossidae or godwit or godwits or monedula or pongidae or atheriniformes or colobinae or lophocebus or atelidae or cottidae or leucopsis or acanthuridae or didelphimorphia or elver or elvers or lapponica or dermoptera or european hake or european hakes or gerbillinae or banteng or hartebeest or hartebeests or hogget or haematopus or anguis fragilis or grey heron or grey herons or blue whiting or blue whitings or furnariidae or macrovipera or esocidae or lapwing or lapwings or mylopharyngodon or wallabia or beloniformes or potoroo or potoroos or athene noctua or pleuronectidae or bushbabies or muscicapidae or alligatoridae or fuligula or bush baby or guineafowl or spoonbill or spoonbills or viverridae or catostomidae or zebrafishes or ibexes or vendace or estrildidae or monotremata or sepiella or ambystomatidae or shelduck or shelducks or treeshrew or treeshrews or hoplobatrachus or pochard or hoolock or hoolocks or lynxes or antilope or antilopes or blackbuck or blackbucks or cricetinae or paramisgurnus or skylark or skylarks or soleidae or allobates or northern wheatear or northern wheatears or pitheciidae or takin or theria or vanellus or galaxiidae or lorisidae or ostralegus or palaeognathae or stone loach or alauda or callitrichinae or caniformia or duttaphrynus or ictaluridae or osteoglossiformes or poultries or curema or ruddy turnstone or ruddy turnstones or sheatfish or sunfishes or centropomidae or hemachatus or platalea or thamnophilidae or song thrush or atherinopsidae or siluridae or tadorna or chroicocephalus or ermine or ermines or gavialis or ruff or tupaiidae or diprotodontia or hyaenidae or antilopinae or crocodylidae or herpestidae or hippopotamidae or northern shoveler or round gobies or cheirogaleidae or indriidae or fundulidae or pythonidae or rhynchocephalia or anodorhynchus or red-backed shrike or red-backed shrikes or triakidae or phalangeridae or aoudad or boreoeutheria or eurasianjay or eurasian jays or feliformia or haplorhini or osteoglossidae or paenungulata or struthioniformes or ferina or sanderling or sanderlings or spheniscidae or cuttlefishes or cygnet or dasycneme or gadwall or gadwalls or pelobates fuscus or wryneck or wrynecks or afrosoricida or culaea or dover sole or dover soles or paralichthyidae or passeridae or osteolaemus or song thrushes or bluethroat or bluethroats or hydrophiidae or megrim or mephitidae or strepsirhini or tomistoma or epidalea or osmeriformes or bush babies or tarsiiform or atelinae or bufotes or eurasian coot or eurasian coots or galagidae or geopelia or philomachus or tubulidentata or bombinatoridae or pelobatidae or tachysurus or ailuridae or woodlark or woodlarks or alcelaphinae or redshank or redshanks or salientia or sand smelt or sand smelts or woodmice or woodmouse or dasyproctidae or eurasian wigeon or eurasian wigeons or garganey or garganeys or lemon sole or lemon soles or common dab or common dabs or graylag or graylags or leucorodia or osphronemidae or bewickii or common moorhen or common moorhens or decapodiformes or gobbler or gobblers or odontophoridae or paddlefishes or eutheria or salmonine or esociformes or eurasian woodcock or eurasian woodcocks or european smelt or european smelts or goldfishes or tenches or tyranni or common chaffinch or common chaffinchs or common redstart or common redstarts or common roach or common roachs or great knot or great knots or potoroidae or alytidae or coregonine or dipteral or leveret or poeciliopsis gracilis or amphiumidae or batrachoidiformes or bighead goby or heteropneustidae or lullula or norway pout or norway pouts or sipunculida or dogfishes or sebastidae or tarsiidae or alethinophidia or common nase or common nases or common sandpiper or common sandpipers or eurasian blackcap or eurasian blackcaps or pterocnemia or syngnathiformes or common chaffinches or eupleridae or octopodiformes or phascolarctidae or scophthalmidae or starry smooth-hound or starry smooth-hounds or whitefishes or cuniculidae or european sprat or european sprats or rosy bitterling or rosy bitterlings or common dace or common daces or lesser weever or lesser weevers or scaldfish or water rail or water rails or alouattinae or centrarchiformes or common whitethroat or common whitethroats or gavialidae or grey gurnard or grey gurnards or lateolabracidae or rheiformes or tubgurnard or tub gurnards or common chiffchaff or common chiffchaffs or garfishes or lesser whitethroat or lesser whitethroats or myoxidae or seabasses or spariformes or umbridae or yellow boxfish or anabantiformes or aotidae or common bleak or common bleaks or common rudd or common rudds or greater pipefish or hapale or nandiniidae or stone loaches or whinchat or whinchats or acanthuriformes or brotula barbata or common ling or common lings or common roaches or cottonrat or cottonrats or douroucoulis or dromaiidae or fitches or fitchew or galaxiiformes or laprine or saimiriinae or solenette or tarsii or tompot blenny or common dragonet or common dragonets or longspined bullhead or longspined bullheads or monotremate or monotremates or pempheriformes or perdicinae or presbytini or smegmamorpha or bighead gobies or carangaria incertae sedis or coiidae or fivebeard rockling or foulmart or foumart or grasskeet or greater pipefishes or ibices or millionfish or muguliformes or norwegian topknot or peewit or red sea sailfin tang or rupicapras or sheatfishes or tompot blennies or twait shad or yellow boxfishes).tw. 14176107

3 medline.st. 29904626

4 2 not 3 8398417

5 1 or 4 40352035

6 "Receptors, CCR5"/ 18466

7 (cc chemokine receptor* 5 or cc-ckr5 or ccr5 or cd195 antigen* or ckr5 receptor*).tw,kf. 22382

8 exp CCR5 Receptor Antagonists/ 8211

9 C-C chemokine receptor type 5.tw,kf. 434

10 (md6p741w8a or maraviroc or selzentry or Celsentri or uk 427,857 or uk-427,857 or uk427,857).tw,kf. 3408

11 Vicriviroc.mp. 728

12 (Aplaviroc or ancriviroc or aplaviroc or leronlimab or mavorixafor).mp. 866

13 INCB009471.mp. 2

14 TBR 652.mp. 37

15 Pro 140.mp. 297

16 HGS004.mp. 15

17 6 or 7 or 8 or 9 or 10 or 11 or 12 or 13 or 14 or 15 or 16 33160

18 5 and 17 23661

19 exp cerebrovascular disorders/ or stroke/ 1287736

20 stroke*.tw,kf. 792358

21 transient isch?em* attack*.tw,kf. 42566

22 brain injur*.tw,kf. 199884

23 ((brain or cereb* or intracereb*) adj2 (isch?em* or infarct* or h?emorrhag*)).tw,kf. 261843

24 or/19-23 1714778

25 18 and 24 373

26 use medall 58

27 exp animal experiment/ or exp animal model/ or exp experimental animal/ or exp transgenic animal/ or exp male animal/ or exp female animal/ or exp juvenile animal/ or animal/ or chordata/ or vertebrate/ or tetrapod/ or exp fish/ or amniote/ or exp amphibia/ or mammal/ or exp reptile/ or exp sauropsid/ or therian/ or exp monotreme/ or placental mammal/ or exp marsupial/ or Euarchontoglires/ or exp Afrotheria/ or exp Boreoeutheria/ or exp Laurasiatheria/ or exp Xenarthra/ or primate/ or exp Dermoptera/ or exp Glires/ or exp Scandentia/ or Haplorhini/ or exp prosimian/ or simian/ or exp tarsiiform/ or Catarrhini/ or exp Platyrrhini/ or ape/ or exp Cercopithecidae/ or hominid/ or exp hylobatidae/ or exp chimpanzee/ or exp gorilla/ or exp orang utan/ or exp cephalopod/ 14616185

28 (rat or rats or animal or animals or mice or "in vivo" or mouse or rabbit or rabbits or murine or pig or pigs or dog or dogs or bovine or fish or vertebrate or vertebrates or cat or cats or rodent or rodents or mammal or mammals or chicken or chickens or monkey or monkeys or sheep or canine or canines or porcine or cattle or bird or birds or hamster or hamsters or primate or primates or cow or cows or chick or horse or horses or avian or avians or calf or swine or swines or xenopus or turkeys or bear or bears or frog or frogs or zebrafish or goat or goats or equine or calves or poultry or macaque or macaques or mole or moles or ovine or lamb or lambs or fishes or diptera or amphibian or amphibians or snake or snakes or ruminant or ruminants or hen or hens or piglet or piglets or feline or felines or simian or simians or laevis or trout or trouts or teleost or teleosts or salmon or salmons or seal or seals or bull or bulls or ewe or ewes or hedgehog or hedgehogs or macaca or macacas or proteus or pigeon or pigeons or bat or bats or duck or ducks or chimpanzee or chimpanzees or baboon or baboons or deer or deers or rana or ranas or carp or carps or heifer or swallow or swallows or lizard or lizards or canis or sow or sows or cynomolgus or quail or quails or reptile or reptiles or turtle or turtles or buffalo or gerbil or gerbils or boar or boars or squirrel or squirrels or oncorhynchus or mus or toad or toads or fowl or fowls or rerio or danio or ara or aras or musculus or tadpole or tadpoles or mulatta or salmo or ram or eagle or eagles or ferret or ferrets or goldfish or catfish or whale or whales or fox or foxes or ape or apes or elephant or elephants or bos or marmoset or marmosets or cod or cods or shark or sharks or wolf or eel or eels or auratus or rattus or zebra or zebras or tilapia or tilapias or gilt or camel or camels or squid or gallus or marsupial or marsupials or vole or voles or fascicularis or ovis or salmonid or salmonids or tiger or tigers or dolphin or dolphins or robin or robins or carpio or opossum or opossums or cyprinus or salamander or salamanders or felis or mink or minks or swan or swans or norvegicus or bufo or torpedo or bass or lamprey or lampreys or sus or python or pythons or tetrapod or tetrapods or shrew or shrews or lion or lions or hog or hogs or songbird or songbirds or oreochromis or starling or starlings or caprine or carassius or owl or owls or newt or newts or papio or scrofa or hare or hares or gorilla or gorillas or flounder or flounders or goose or herring or herrings or therian or buffaloes or canary or sparrow or sparrows or microtus or octopus or troglodytes or tuna or amphibia or chinchilla or chinchillas or ide or oryzias or cervus or kangaroo or kangaroos or armadillo or armadillos or callithrix or "pan troglodytes" or saimiri or cichlid or cichlids or donkey or donkeys or bream or char or chars or finch or raccoon or raccoons or bothrops or anguilla or perch or cricetus or seabird or seabirds or buck or bucks or naja or coturnix or salmonids or geese or minnow or minnows or raptor or raptors or merione or meriones or rodentia or elaphus or amniote or amniotes or elasmobranch or emu or emus or peromyscus or hominid or hominids or bubalus or crotalus or gull or gulls or anas or anura or lemur or lemurs or crow or crows or camelus or gibbon or gibbons or waterfowl or parrot or parrots or eels or cob or stickleback or sticklebacks or columba or mesocricetus or ambystoma or raven or ravens or gadus or penguin or penguins or orangutan or orangutans or sturgeon or sturgeons or cuniculus or aves or virginianus or cephalopod or cephalopods or cebus or sparus or tortoise or tortoises or guttata or morhua or unguiculatus or dogfish or vulpes or mallard or mallards or apodemus or alligator or alligators or oryctolagus or llama or llamas or reindeer or mustela or duckling or ducklings or wolves or sander or amazona or zebu or badger or badgers or dove or doves or ictalurus or capra or capras or equus or camelid or camelids or poecilia or mule or mules or perciformes or salvelinus or labrax or cyprinidae or ariidae or crocodile or crocodiles or fundulus or dicentrarchus or clarias or cercopithecus or chiroptera or alpaca or alpacas or pike or pikes or paralichthys or puma or pumas or didelphis or pisces or macropus or triturus or bison or bisons or epinephelus or gasterosteus or panthera or acipenser or mackerel or mackerels or tamarin or tamarins or ostrich or anolis or vervet or vervets or wallaby or glareolus or beaver or beavers or dromedary or catus or killifish or pimephales or promelas or aotus or phoca or panda or pandas or porpoise or porpoises or myotis or yak or yaks or agkistrodon or vipera or otter or otters or turbot or turbots or squamate or carnivora or mullet or mullets or hawk or hawks or taeniopygia or seahorse or seahorses or "poecilia reticulata" or falcon or falcons or prosimian or prosimians or parus or perca or fingerling or fingerlings or antelope or antelopes or tupaia or passeriformes or sepia or saguinus or coyote or coyotes or pongo or meleagris or reptilia or lepus or psittacine or hagfish or warbler or warblers or "russell s viper" or "russell s vipers" or smolt or smolts or budgerigar or sardine or sardines or cavia or cavias or hyla or pleurodeles or siluriformes or "great tit" or "great tits" or guppy or bonobo or bonobos or rutilus or trichosurus or muridae or phodopus or channa or squalus or lynx or sturnus or petromyzon or vitulina or monodelphis or cuttlefish or adder or adders or lepomis or canaria or gambusia or guppies).tw. 14111285

29 (xiphophorus or flatfish or koala or koalas or labeo or stingray or stingrays or chelonia or lampetra or spermophilus or crocodilian or "passer domesticus" or sciurus or artiodactyla or ranidae or corvus or necturus or platypus or canaries or bovid or lagopus or trimeresurus or gariepinus or marten or martens or drosophilidae or mugil or sunfish or porcellus or cypriniformes or alouatta or scophthalmus or anser or electrophorus or putorius or iguana or iguanas or lama or lamas or takifugu or circus or eptesicus or flycatcher or galago or galagos or trachemys or lungfish or characiformes or shorebird or shorebirds or giraffe or giraffes or micropterus or scyliorhinus or cichlidae or loligo or porcupine or porcupines or chub or chubs or solea or pleuronectes or hylidae or viperidae or echis or sorex or anchovy or lagomorph or ostriches or vulture or vultures or whitefish or araneus or jird or jirds or tern or esox or drake or drakes or elapidae or gallopavo or chordata or myodes or caretta or serinus or grouse or misgurnus or meles or blackbird or blackbirds or coregonus or bobwhite or bobwhites or heteropneustes or mammoth or mammoths or turdus or rhinella or ateles or characidae or clupea or bungarus or brill or "struthio camelus" or sloth or sloths or pteropus or sculpin or anthropoids or pollock or pollocks or morone or "pan paniscus" or litoria or chipmunk or chipmunks or balaenoptera or marmota or melopsittacus or hyrax or lemming or lemmings or halibut or hylobates or lates or caiman or caimans or sigmodon or stenella or barbel or barbels or sterna or parakeet or parakeets or phocoena or leptodactylus or canidae or buteo or harengus or gopher or gophers or marmot or marmots or gosling or goslings or platichthys or gar or gars or sebastes or marsupialia or notophthalmus or gazelle or gazelles or insectivora or paridae or felidae or russula or galliformes or bombina or colobus or echidna or echidnas or seabass or syncerus or plaice or "blue tit" or "blue tits" or pagrus or catfishes or cetacea or barbus or cygnus or ficedula or chamois or colubridae or perches or coelacanth or fitch or urodela or cynops or martes or halichoerus or aix or salmonidae or leuciscus or magpie or magpies or silurus or whiting or whitings or anseriformes or colinus or rhea or chlorocebus or octodon or acinonyx or mouflon or mouflons or ibex or tetraodon or bufonidae or equidae or jackal or cephalopoda or dendroaspis or glama or muskrat or muskrats or sable or sables or wildebeest or streptopelia or albifrons or vespertilionidae or woodpecker or woodpeckers or muntjac or muntjacs or archosaur or branta or cricetulus or megalobrama or poeciliidae or desmodus or snakehead or snakeheads or tench or teal or teals or bandicoot or bandicoots or apteronotus or phyllostomidae or crocidura or buzzard or buzzards or larimichthys or cercocebus or pipistrellus or erithacus or impala or impalas or rousettus or haddock or haddocks or tinca or ratite or calidris or cynoglossus or hypophthalmichthys or bullock or bullocks or dromedaries or alectoris or filly or salamandra or cingulata or bitis or grus or ammodytes or macaw or macaws or hypoleuca or sapajus or cyprinodontiformes or hippopotamus or pelophylax or capybara or capybaras or weasel or weasels or cairina or cynomys or lutra or cockatoo or cockatoos or lachesis or lagomorpha or rupicapra or daboia or "orang utan" or "orang utans" or platyrrhini or charadriiformes or micrurus or psittaciformes or spalax or loris or mustelidae or sylvilagus or vitticeps or cockatiel or mustelus or cottus or erythrocebus or dipodomys or platessa or callicebus or loricariidae or catostomus or cuneata or cyanistes or cyprinodon or sigmodontinae or elasmobranchii or trichechus or sauropsid or xenarthra or dormouse or perissodactyla or nautilus or cirrhinus or gulo or gulos or tragelaphus or merula or numida or sciaenidae or cerastes or sciuridae or gibbosus or octopuses or eland or elands or phyllomedusa or pogona or walrus or agamidae or leptodactylidae or ridibundus or leontopithecus or anteater or anteaters or pelodiscus or cebidae or columbianus or "pelteobagrus fulvidraco" or hominoidea or mandrillus or "zonotrichia leucophrys" or agama or gobiocypris or "bearded dragon" or "bearded dragons" or sarotherodon or talpa or discoglossus or hagfishes or sphenodon or gudgeon or amphiuma or aythya or tenrec or tenrec or hominidae or risoria or salamandridae or camelidae or columbiformes or latimeria or plover or plovers or afrotheria or "falco sparverius" or polecat or polecats or crotalinae or salvadora or tarsier or lucioperca or anchovies or lungfishes or terrapin or "dromaius novaehollandiae" or lateolabrax or eigenmannia or pelamis or theropithecus or murinae or gander or gymnotus or pseudacris or gymnophiona or gymnotiformes or laticauda or falconiformes or dugong or dugongs or pintail or pintails or rook or rooks or lasiurus or catshark or catsharks or micropogonias or "red junglefowl" or paddlefish or ophiophagus or hollandicus or nymphicus or pimelodidae or aepyceros or cobitidae or strigiformes or cobitis or dormice or alytes or calloselasma or guanaco or guanacos or phasianidae or "round goby" or trichogaster or catarrhini or eelpout or eelpouts or galaxias or gaur or pungitius or suslik or susliks or flatfishes or percidae or caprinae or todarodes or osmerus or ameiurus or anthropoidea or "castor canadensis" or pouting or poutings or tetraodontiformes or arvicolinae or siamang or siamangs or "castor fiber" or nomascus or "red knot" or "red knots" or syngnathidae or iguanidae or eretmochelys or ursidae or callimico or columbidae or microhylidae or anaxyrus or menidia or pipistrelle or greylag or pipidae or scandentia or bowfin or bowfins or dendrobatidae or zenaida or bushbaby or harrier or harriers or macropodidae or pygerythrus or clupeidae or odorrana or corvidae or jerboa or jerboas or canutus or hylobatidae or clupeiformes or "great cormorant" or "great cormorants" or scorpaeniformes or chondrostean or garfish or proboscidea or psetta or diapsid or serotinus or tetrao or walruses or carcharhiniformes or leucoraja or pumpkinseed or dosidicus or acipenseriformes or daubentonii or emberizidae or gadiformes or hyraxes or stizostedion or wolverine or wolverines or lissotriton or acanthurus or centrarchidae or gloydius or laurasiatheria or limosa or psittacula or leporidae or proteidae or zander or zanders or arapaima or bagridae or cyprinodontidae or mithun or pandion or jackdaw or jackdaws or procyonidae or carus or jaculus or salmoniformes or "common sole" or "common soles" or protobothrops or calamita or brachyteles or trionyx or turdidae or boidae or luscinia or pugnax or euarchontoglires or saithe or saithes or symphalangus or aardvark).tw. 272058

30 (aardvarks or oystercatcher or oystercatchers or arius or corydoras or poacher or poachers or aurochs or cebuella or crecca or lemuridae or sirenia or lemmus or perdix or glires or lepidosaur or muskox or deinagkistrodon or pholidota or holocephali or cercopithecinae or clariidae or agapornis or doryteuthis or tyrannidae or dicroglossidae or godwit or godwits or monedula or pongidae or atheriniformes or colobinae or lophocebus or atelidae or cottidae or leucopsis or acanthuridae or didelphimorphia or elver or elvers or lapponica or dermoptera or "european hake" or "european hakes" or gerbillinae or banteng or hartebeest or hartebeests or hogget or haematopus or "anguis fragilis" or "grey heron" or "grey herons" or "blue whiting" or "blue whitings" or furnariidae or macrovipera or esocidae or lapwing or lapwings or mylopharyngodon or wallabia or beloniformes or potoroo or potoroos or "athene noctua" or pleuronectidae or bushbabies or muscicapidae or alligatoridae or fuligula or "bush baby" or guineafowl or spoonbill or spoonbills or viverridae or catostomidae or zebrafishes or ibexes or vendace or estrildidae or monotremata or sepiella or ambystomatidae or shelduck or shelducks or treeshrew or treeshrews or hoplobatrachus or pochard or hoolock or hoolocks or lynxes or antilope or antilopes or blackbuck or blackbucks or cricetinae or paramisgurnus or skylark or skylarks or soleidae or allobates or "northern wheatear" or "northern wheatears" or pitheciidae or takin or theria or vanellus or galaxiidae or lorisidae or ostralegus or palaeognathae or "stone loach" or alauda or callitrichinae or caniformia or duttaphrynus or ictaluridae or osteoglossiformes or poultries or curema or "ruddy turnstone" or "ruddy turnstones" or sheatfish or sunfishes or centropomidae or hemachatus or platalea or thamnophilidae or "song thrush" or atherinopsidae or siluridae or tadorna or chroicocephalus or ermine or ermines or gavialis or ruff or tupaiidae or diprotodontia or hyaenidae or antilopinae or crocodylidae or herpestidae or hippopotamidae or "northern shoveler" or "round gobies" or cheirogaleidae or indriidae or fundulidae or pythonidae or rhynchocephalia or anodorhynchus or "red-backed shrike" or "red-backed shrikes" or triakidae or phalangeridae or aoudad or boreoeutheria or "eurasian jay" or "eurasian jays" or feliformia or haplorhini or osteoglossidae or paenungulata or struthioniformes or ferina or sanderling or sanderlings or spheniscidae or cuttlefishes or cygnet or dasycneme or gadwall or gadwalls or "pelobates fuscus" or wryneck or wrynecks or afrosoricida or culaea or "dover sole" or "dover soles" or paralichthyidae or passeridae or osteolaemus or "song thrushes" or bluethroat or bluethroats or hydrophiidae or megrim or mephitidae or strepsirhini or tomistoma or epidalea or osmeriformes or "bush babies" or tarsiiform or atelinae or bufotes or "eurasian coot" or "eurasian coots" or galagidae or geopelia or philomachus or tubulidentata or bombinatoridae or pelobatidae or tachysurus or ailuridae or woodlark or woodlarks or alcelaphinae or redshank or redshanks or salientia or "sand smelt" or "sand smelts" or woodmice or woodmouse or dasyproctidae or "eurasian wigeon" or "eurasian wigeons" or garganey or garganeys or "lemon sole" or "lemon soles" or "common dab" or "common dabs" or graylag or graylags or leucorodia or osphronemidae or bewickii or "common moorhen" or "common moorhens" or decapodiformes or gobbler or gobblers or odontophoridae or paddlefishes or eutheria or salmonine or esociformes or "eurasian woodcock" or "eurasian woodcocks" or "european smelt" or "european smelts" or goldfishes or tenches or tyranni or "common chaffinch" or "common chaffinchs" or "common redstart" or "common redstarts" or "common roach" or "common roachs" or "great knot" or "great knots" or potoroidae or alytidae or coregonine or dipteral or leveret or "poeciliopsis gracilis" or amphiumidae or batrachoidiformes or "bighead goby" or heteropneustidae or lullula or "norway pout" or "norway pouts" or sipunculida or dogfishes or sebastidae or tarsiidae or alethinophidia or "common nase" or "common nases" or "common sandpiper" or "common sandpipers" or "eurasian blackcap" or "eurasian blackcaps" or pterocnemia or syngnathiformes or "common chaffinches" or eupleridae or octopodiformes or phascolarctidae or scophthalmidae or "starry smooth-hound" or "starry smooth-hounds" or whitefishes or cuniculidae or "european sprat" or "european sprats" or "rosy bitterling" or "rosy bitterlings" or "common dace" or "common daces" or "lesser weever" or "lesser weevers" or scaldfish or "water rail" or "water rails" or alouattinae or centrarchiformes or "common whitethroat" or "common whitethroats" or gavialidae or "grey gurnard" or "grey gurnards" or lateolabracidae or rheiformes or "tub gurnard" or "tub gurnards" or "common chiffchaff" or "common chiffchaffs" or garfishes or "lesser whitethroat" or "lesser whitethroats" or myoxidae or seabasses or spariformes or umbridae or "yellow boxfish" or anabantiformes or aotidae or "common bleak" or "common bleaks" or "common rudd" or "common rudds" or "greater pipefish" or hapale or nandiniidae or "stone loaches" or whinchat or whinchats or acanthuriformes or "brotula barbata" or "common ling" or "common lings" or "common roaches" or cottonrat or cottonrats or douroucoulis or dromaiidae or fitches or fitchew or galaxiiformes or laprine or saimiriinae or solenette or tarsii or "tompot blenny" or "common dragonet" or "common dragonets" or "longspined bullhead" or "longspined bullheads" or monotremate or monotremates or pempheriformes or perdicinae or presbytini or smegmamorpha or "bighead gobies" or "carangaria incertae sedis" or coiidae or "fivebeard rockling" or foulmart or foumart or grasskeet or "greater pipefishes" or ibices or millionfish or muguliformes or "norwegian topknot" or peewit or "red sea sailfin tang" or rupicapras or sheatfishes or "tompot blennies" or "twait shad" or "yellow boxfishes").tw. 21723

31 or/27-30 18002118

32 exp chemokine receptor CCR5 antagonist/ 6682

33 (cc chemokine receptor* 5 or cc-ckr5 or ccr5 or cd195 antigen* or ckr5 receptor* or C-C chemokine receptor type 5).tw. 22155

34 (md6p741w8a or maraviroc or selzentry or Celsentri or uk 427,857 or uk-427,857 or uk427,857).tw. 3329

35 Vicriviroc.mp. 728

36 (Aplaviroc or ancriviroc or aplaviroc or leronlimab or mavorixafor).mp. 866

37 INCB009471.mp. 2

38 TBR 652.mp. 37

39 Pro 140.mp. 297

40 HGS004.mp. 15

41 chemokine receptor CCR5/ 12698

42 or/32-41 31956

43 31 and 42 10275

44 exp cerebrovascular disease/ or exp cerebrovascular accident/ 1207681

45 stroke*.tw. 771477

46 ((brain or cereb* or intracereb*) adj2 (isch?em* or infarct* or h?emorrhag*)).tw. 247502

47 (transient isch?em* attack* or brain injur*).tw. 231020

48 44 or 45 or 46 or 47 1675247

49 43 and 48 172

50 use emczd 118

51 26 or 50 176

**\*\*\*\*\*\*\*\*\*\*\*\*\*\*\*\*\*\*\*\*\*\*\*\*\*\*\*\*\*\*\*\*\*\*\*\*\*\*\*\*\*\*\*\*\*\*\*\*\*\*\*\*\*\*\*\*\*\*\*\*\*\*\*\*\*\*\*\*\*\*\*\*\*\*\*\*\*\***

**Web of Science – October 25, 2022**

\# Web of Science Search Strategy (v0.1)

\# Entitlements:

- WOS.SSCI: 1900 to 2022

- WOS.AHCI: 1975 to 2022

- WOS.ISTP: 1990 to 2022

- WOS.ESCI: 2005 to 2022

- WOS.SCI: 1900 to 2022

- WOS.ISSHP: 1990 to 2022

\# Searches:

1: CCR5 (Topic) Results: 11678

2: TS=(cc chemokine receptor* 5) OR TS=(C-C chemokine receptor type 5) Results: 1796

3: TS=(cc-ckr5) Results: 9

4: TS=(Vicriviroc) Results: 158

5: TS=(((md6p741w8a OR maraviroc OR selzentry OR Celsentri))) Results: 1542

6: ALL=((Aplaviroc or ancriviroc or aplaviroc or leronlimab or mavorixafor)) Results: 97

7: (((TS=(INCB009471)) OR TS=(TBR 652)) OR TS=(Pro 140)) OR TS=(HGS004)

Results: 972

8: #7 OR #6 OR #5 OR #4 OR #3 OR #2 OR #1 Results: 14241

9: (TS=(stroke*)) OR TS=("transient isch?em* attack*") Results: 430668

10: TS=((brain or cereb* or intracereb*) NEAR/2 ischem*) Results: 77919

11: TS=(((brain or cereb* or intracereb*) NEAR/2 ischaem*)) Results: 5389

12: TS=(((brain or cereb* or intracereb*) NEAR/2 hemorrhag*)) Results: 33959

13: TS=(((brain or cereb* or intracereb*) NEAR/2 haemorrhag*)) Results: 4867

14: TS="cerebrovascular disorder*" Results: 3993

15: TS="brain injur*" Results: 121601

16: #15 OR #14 OR #13 OR #12 OR #11 OR #10 OR #9 Results: 591495

17: #8 AND #16 Results: 136

18: TS=(rat OR rats OR animal OR animals OR mice OR "in vivo" OR mouse OR rabbit OR rabbits OR murine OR pig OR pigs OR dog OR dogs OR bovine OR fish OR vertebrate OR vertebrates OR cat OR cats OR rodent OR rodents OR mammal OR mammals OR chicken OR chickens OR monkey OR monkeys OR sheep OR canine OR canines OR porcine OR cattle OR bird OR birds OR hamster OR hamsters OR primate OR primates OR cow OR cows OR chick OR horse OR horses OR avian OR avians OR calf OR swine OR swines OR xenopus OR turkeys OR bear OR bears OR frog OR frogs OR zebrafish OR goat OR goats OR equine OR calves OR poultry OR macaque OR macaques OR mole OR moles OR ovine OR lamb OR lambs OR fishes OR diptera OR amphibian OR amphibians OR snake OR snakes OR ruminant OR ruminants OR hen OR hens OR piglet OR piglets OR feline OR felines OR simian OR simians OR laevis OR trout OR trouts OR teleost OR teleosts OR salmon OR salmons OR seal OR seals OR bull OR bulls OR ewe OR ewes OR hedgehog OR hedgehogs OR macaca OR macacas OR proteus OR pigeon ORpigeons OR bat OR bats OR duck OR ducks OR chimpanzee OR chimpanzees OR baboon OR baboons OR deer OR rana OR ranas OR carp OR carps OR heifer OR swallow OR swallows OR lizard OR lizards OR canis OR sow OR sows OR cynomolgus OR quail OR quails OR reptile OR reptiles OR turtle OR turtles OR buffalo OR gerbil OR gerbils OR boar OR boars OR squirrel OR squirrels OR oncorhynchus OR mus OR toad OR toads OR fowl OR fowls OR rerio OR danio OR ara OR aras OR musculus OR tadpole OR tadpoles OR mulatta OR salmo OR ram OR eagle OR eagles OR ferret OR ferrets OR goldfish OR catfish OR whale OR whales OR fox OR foxes OR ape OR apes OR elephant OR elephants OR bos OR marmoset OR marmosets OR cod OR cods OR shark OR sharks OR wolf OR eel OR eels OR auratus OR rattus OR zebra OR zebras OR tilapia OR tilapias OR gilt OR camel OR camels OR squid OR gallus OR marsupial OR marsupials OR vole OR voles OR fascicularis OR ovis OR salmonid OR salmonids OR tiger OR tigers OR dolphin OR dolphins OR robin OR robins OR carpio OR opossumOR opossums OR cyprinus OR salamander OR salamanders OR felis OR mink OR minks OR swan OR swans OR norvegicus OR bufo OR torpedo OR bass OR lamprey OR lampreys OR sus OR python OR pythons OR tetrapod OR tetrapods OR shrew OR shrews OR lion OR lions OR hogOR hogs OR songbird OR songbirds OR oreochromis OR starling OR starlings OR caprine OR carassius OR owl OR owls OR newt OR newts OR papio OR scrofa OR hare OR hares OR gorilla OR gorillas OR flounder OR flounders OR goose OR herring OR herrings OR therianOR buffaloes OR canary OR sparrow OR sparrows OR microtus OR octopus OR troglodytes OR tuna OR amphibia OR chinchilla OR chinchillas OR ide OR oryzias OR cervus OR kangaroo OR kangaroos OR armadillo OR armadillos OR callithrix OR "pan troglodytes" OR saimiri OR cichlid OR cichlids OR donkey OR donkeys OR bream OR char OR chars OR finch OR raccoon OR raccoons OR bothrops OR anguilla OR perch OR cricetus OR seabird OR seabirds OR buck OR bucks OR naja OR coturnix OR salmonids OR geese OR minnow OR minnows ORraptor OR raptors OR merione OR meriones OR rodentia OR elaphus OR amniote OR amniotes OR elasmobranch OR emu OR emus OR peromyscus OR hominid OR hominids OR bubalus OR crotalus OR gull OR gulls OR anas OR anura OR lemur OR lemurs OR crow OR crows OR camelus OR gibbon OR gibbons OR waterfowl OR parrot OR parrots OR eels OR cob OR stickleback OR sticklebacks OR columba OR mesocricetus OR ambystoma OR raven OR ravens OR gadus OR penguin OR penguins OR orangutan OR orangutans OR sturgeon OR sturgeons OR cuniculus OR aves OR virginianus OR cephalopod OR cephalopods OR cebus OR sparus OR tortoise OR tortoises OR guttata OR morhua OR unguiculatus OR dogfish OR vulpes OR mallard OR mallards OR apodemus OR alligator OR alligators OR oryctolagus OR llama OR llamas OR reindeer OR mustela OR duckling OR ducklings OR wolves OR sander OR amazona OR zebu OR badger OR badgers OR dove OR doves OR ictalurus OR capra OR capras OR equus OR camelid OR camelids OR poecilia OR mule OR mules OR perciformes OR salvelinus OR labrax OR cyprinidae OR ariidae OR crocodile OR crocodiles OR fundulus OR dicentrarchus OR clarias OR cercopithecus OR chiroptera OR alpaca OR alpacas OR pike OR pikes OR paralichthys OR puma OR pumas OR didelphis OR pisces OR macropus OR triturus OR bison OR bisons OR epinephelus OR gasterosteus OR panthera OR acipenser OR mackerel OR mackerels OR tamarin OR tamarins OR ostrich OR anolis OR vervet OR vervets OR wallaby OR glareolus OR beaver OR beavers OR dromedary OR catus OR killifish OR pimephales OR promelas OR aotus OR phoca OR panda OR pandas OR porpoise OR porpoises OR myotis OR yak OR yaks OR agkistrodon OR vipera OR otter OR otters OR turbot OR turbots OR squamate OR carnivora OR mullet OR mullets OR hawk OR hawks OR taeniopygia OR seahorse OR seahorses OR "poecilia reticulata" OR falcon OR falcons OR prosimian OR prosimians OR parus OR perca OR fingerling OR fingerlings OR antelope OR antelopes OR tupaia OR passeriformes OR sepia OR saguinus OR coyote OR coyotes OR pongo OR meleagris OR reptilia OR lepus OR psittacine OR hagfish OR warbler OR warblers OR "russell s viper" OR "russell s vipers" OR smolt OR smolts OR budgerigar OR sardine OR sardines OR cavia OR cavias OR hyla OR pleurodeles OR siluriformes OR "great tit" OR "great tits" OR guppy OR bonobo OR bonobos OR rutilus OR trichosurus OR muridae OR phodopus OR channa OR squalus OR lynx OR sturnus OR petromyzon OR vitulina OR monodelphis OR cuttlefish OR adder OR adders OR lepomis OR canaria OR gambusia OR guppies OR xiphophorus OR flatfish OR koala ORkoalas OR labeo OR stingray OR stingrays OR chelonia OR lampetra OR spermophilus OR crocodilian OR "passer domesticus" OR sciurus OR artiodactyla OR ranidae OR corvus OR necturus OR platypus OR canaries OR bovid OR lagopus OR trimeresurus OR gariepinus ORmarten OR martens OR drosophilidae OR mugil OR sunfish OR porcellus OR cypriniformes OR alouatta OR scophthalmus OR anser OR electrophorus OR putorius OR iguana OR iguanas OR lama OR lamas OR takifugu OR circus OR eptesicus OR flycatcher OR galago OR galagos OR trachemys OR lungfish OR characiformes OR shorebird OR shorebirds OR giraffe OR giraffes OR micropterus OR scyliorhinus OR cichlidae OR loligo OR porcupine OR porcupines OR chub OR chubs OR solea OR pleuronectes OR hylidae OR viperidae OR echis OR sorex OR anchovy OR lagomorph OR ostriches OR vulture OR vultures OR whitefish OR araneus OR jird OR jirds OR tern OR esox OR drake OR drakes OR elapidae OR gallopavo OR chordata OR myodes OR caretta OR serinus OR grouse OR misgurnus OR meles OR blackbird OR blackbirds OR coregonus OR bobwhite OR bobwhites OR heteropneustes OR mammoth OR mammoths OR turdus OR rhinella OR ateles OR characidae OR clupea OR bungarus OR brill OR "struthio camelus" OR sloth OR sloths OR pteropus OR sculpin OR anthropoids OR pollock OR pollocks OR morone OR "pan paniscus" OR litoria OR chipmunk OR chipmunks OR balaenoptera OR marmota OR melopsittacus OR hyrax OR lemming OR lemmings OR halibut OR hylobates OR lates OR caiman OR caimans OR sigmodon OR stenella OR barbel OR barbels ORsterna OR parakeet OR parakeets OR phocoena OR leptodactylus OR canidae OR buteo OR harengus OR gopher OR gophers OR marmot OR marmots OR gosling OR goslings OR platichthys OR gar OR gars OR sebastes OR marsupialia OR notophthalmus OR gazelle OR gazelles OR insectivora OR paridae OR felidae OR russula OR galliformes OR bombina OR colobus OR echidna OR echidnas OR seabass OR syncerus OR plaice OR "blue tit" OR "blue tits" OR pagrus OR catfishes OR cetacea OR barbus OR cygnus OR ficedula OR chamois OR colubridae OR perches OR coelacanth OR fitch OR urodela OR cynops OR martes OR halichoerus OR aix OR salmonidae OR leuciscus OR magpie OR magpies OR silurus OR whiting OR whitings OR anseriformes OR colinus OR rhea OR chlorocebus OR octodon OR acinonyx OR mouflon OR mouflons OR ibex OR tetraodon OR bufonidae OR equidae OR jackal OR cephalopoda OR dendroaspis OR glama OR muskrat OR muskrats OR sable OR sables OR wildebeest OR streptopelia OR albifrons OR vespertilionidae OR woodpecker OR woodpeckers OR muntjac OR muntjacs OR archosaur OR branta OR cricetulus OR megalobrama OR poeciliidae OR desmodus OR snakehead OR snakeheads OR tench OR teal OR teals OR bandicoot OR bandicoots OR apteronotus OR phyllostomidae OR crocidura OR buzzard OR buzzards OR larimichthys OR cercocebus OR pipistrellus OR erithacus OR impala OR impalas OR rousettus OR haddock OR haddocks OR tinca OR ratite OR calidris OR cynoglossus OR hypophthalmichthys OR bullock OR bullocks OR dromedaries OR alectoris OR filly OR salamandra OR cingulata OR bitis OR grus OR ammodytes OR macaw OR macaws OR hypoleuca OR sapajus OR cyprinodontiformes OR hippopotamus OR pelophylax OR capybara OR capybaras OR weasel OR weasels OR cairina OR cynomys OR lutra OR cockatoo OR cockatoos OR lachesis OR lagomorpha OR rupicapra OR daboia OR "orang utan" OR "orang utans" OR platyrrhini OR charadriiformes OR micrurus OR psittaciformes OR spalax OR loris OR mustelidae OR sylvilagus OR vitticeps OR cockatiel OR mustelus OR cottus OR erythrocebus OR dipodomys OR platessa OR callicebus OR loricariidae OR catostomus OR cuneata OR cyanistes OR cyprinodon OR sigmodontinae OR elasmobranchii OR trichechus OR sauropsid OR xenarthra OR dormouse OR perissodactyla OR nautilus OR cirrhinus OR gulo OR gulos OR tragelaphus OR merula OR numidaOR sciaenidae OR cerastes OR sciuridae OR gibbosus OR octopuses OR eland OR elands OR phyllomedusa OR pogona OR walrus OR agamidae OR leptodactylidae OR ridibundus OR leontopithecus OR anteater OR anteaters OR pelodiscus OR cebidae OR columbianus OR "pelteobagrus fulvidraco" OR hominoidea OR mandrillus OR "zonotrichia leucophrys" OR agama OR gobiocypris OR "bearded dragon" OR "bearded dragons" OR sarotherodon OR talpa OR discoglossus OR hagfishes OR sphenodon OR gudgeon OR amphiuma OR aythya OR tenrec OR tenrec OR hominidae OR risoria OR salamandridae OR camelidae OR columbiformes OR latimeria OR plover OR plovers OR afrotheria OR "falco sparverius" OR polecat OR polecats OR crotalinae OR salvadora OR tarsier OR lucioperca OR anchovies OR lungfishes OR terrapin OR "dromaius novaehollandiae" OR lateolabrax OR eigenmannia OR pelamis OR theropithecus OR murinae OR gander OR gymnotus OR pseudacris OR gymnophiona OR gymnotiformes OR laticauda OR falconiformes OR dugong OR dugongs OR pintail OR pintails OR rook ORrooks OR lasiurus OR catshark OR catsharks OR micropogonias OR "red junglefowl" OR paddlefish OR eutheria OR ophiophagus OR hollandicus OR nymphicus OR pimelodidae OR aepyceros OR cobitidae OR strigiformes OR cobitis OR dormice OR alytes OR calloselasma OR guanaco OR guanacos OR phasianidae OR "round goby" OR trichogaster OR catarrhini OR eelpout OR eelpouts OR galaxias OR gaur OR pungitius OR suslik OR susliks OR flatfishes OR percidae OR caprinae OR todarodes OR osmerus OR ameiurus OR anthropoidea OR "castor canadensis" OR pouting OR poutings OR tetraodontiformes OR arvicolinae OR siamang OR siamangs OR "castor fiber" OR nomascus OR "red knot" OR "red knots" OR syngnathidae OR iguanidae OR eretmochelys OR ursidae OR callimico OR columbidae OR microhylidaeOR anaxyrus OR menidia OR pipistrelle OR greylag OR pipidae OR scandentia OR bowfin OR bowfins OR dendrobatidae OR zenaida OR bushbaby OR harrier OR harriers OR macropodidae OR pygerythrus OR clupeidae OR odorrana OR corvidae OR jerboa OR jerboas OR canutus OR hylobatidae OR clupeiformes OR "great cormorant" OR "great cormorants" OR scorpaeniformes OR chondrostean OR garfish OR proboscidea OR psetta OR diapsid OR serotinus OR tetrao OR walruses OR carcharhiniformes OR leucoraja OR pumpkinseed OR dosidicus OR acipenseriformes OR daubentonii OR emberizidae OR gadiformes OR hyraxes OR stizostedion OR wolverine OR wolverines OR lissotriton OR acanthurus OR centrarchidae OR gloydius OR laurasiatheria OR limosa OR psittacula OR leporidae OR proteidae OR zander ORzanders OR arapaima OR bagridae OR cyprinodontidae OR mithun OR pandion OR jackdaw OR jackdaws OR procyonidae OR carus OR jaculus OR salmoniformes OR "common sole" OR "common soles" OR protobothrops OR calamita OR brachyteles OR trionyx OR turdidae OR boidae OR luscinia OR pugnax OR euarchontoglires OR saithe OR saithes OR symphalangus OR aardvark OR aardvarks OR oystercatcher OR oystercatchers OR arius OR corydoras OR poacher OR poachers OR aurochs OR cebuella OR crecca OR lemuridae OR sirenia OR lemmus OR perdix OR glires OR lepidosaur OR muskox OR deinagkistrodon OR pholidota OR holocephali OR cercopithecinae OR clariidae OR agapornis OR doryteuthis OR tyrannidae OR dicroglossidae OR godwit OR godwits OR monedula OR pongidae OR atheriniformes OR colobinae OR lophocebus OR atelidae OR cottidae OR leucopsis OR acanthuridae OR didelphimorphia OR elver OR elvers OR lapponica OR dermoptera OR "european hake" OR "european hakes" OR gerbillinae OR banteng OR hartebeest OR hartebeests OR hogget OR haematopus OR "anguis fragilis" OR "grey heron" OR "grey herons" OR "blue whiting" OR "blue whitings" OR furnariidae OR macrovipera OR esocidae OR lapwing OR lapwings OR mylopharyngodon OR wallabia OR beloniformes OR potoroo OR potoroos OR "athene noctua" OR pleuronectidae OR bushbabies OR muscicapidae OR alligatoridae OR fuligula OR "bush baby" OR guineafowl OR spoonbill OR spoonbills OR viverridae OR catostomidae OR zebrafishes OR ibexes OR vendace OR estrildidae OR monotremata OR sepiella OR ambystomatidae OR shelduck OR shelducks OR treeshrew OR treeshrews OR hoplobatrachus OR pochard OR hoolock OR hoolocks OR lynxes OR antilope OR antilopes OR blackbuck OR blackbucks OR cricetinae OR paramisgurnus OR skylark OR skylarks OR soleidae OR allobates OR "northern wheatear" OR "northern wheatears" OR pitheciidae OR takin OR theria OR vanellus OR galaxiidae OR lorisidae OR ostralegus OR palaeognathae OR "stone loach" OR alauda OR callitrichinae OR caniformia OR duttaphrynus OR ictaluridae OR osteoglossiformes OR poultries OR curema OR "ruddy turnstone" OR "ruddy turnstones" OR sheatfish OR sunfishes OR centropomidae OR hemachatus OR platalea OR thamnophilidae OR "song thrush" OR atherinopsidae OR siluridae OR tadorna OR chroicocephalus OR ermine OR ermines OR gavialis OR ruffe OR tupaiidae OR diprotodontia OR hyaenidae OR antilopinae OR crocodylidae OR herpestidae OR hippopotamidae OR "northern shoveler" OR "round gobies" OR cheirogaleidae OR indriidae OR fundulidae OR pythonidae OR rhynchocephalia OR anodorhynchus OR "red-backed shrike" OR "red-backed shrikes" OR triakidae OR phalangeridae OR aoudad OR boreoeutheria OR "eurasian jay" OR "eurasian jays" OR feliformia OR haplorhini OR osteoglossidae OR paenungulata OR struthioniformes OR ferina OR sanderling OR sanderlings OR spheniscidae OR cuttlefishes OR cygnet OR dasycneme OR gadwall OR gadwalls OR "pelobates fuscus" OR wryneck OR wrynecks OR afrosoricida OR culaea OR "dover sole" OR "dover soles" OR paralichthyidae OR passeridae OR osteolaemus OR "song thrushes" OR bluethroat OR bluethroats OR hydrophiidae OR megrim OR mephitidae OR strepsirhini OR tomistoma OR epidalea OR osmeriformes OR "bush babies" OR tarsiiform OR atelinae OR bufotes OR "eurasian coot" OR "eurasian coots" OR galagidae OR geopelia OR philomachus OR tubulidentata OR bombinatoridae OR pelobatidae OR tachysurus OR ailuridae OR woodlark OR woodlarks OR alcelaphinae OR redshank OR redshanks OR salientia OR "sand smelt" OR "sand smelts" OR woodmice OR woodmouse OR dasyproctidae OR "eurasian wigeon" OR "eurasian wigeons" OR garganey OR garganeys OR "lemon sole" OR "lemon soles" OR "common dab" OR "common dabs" OR graylag OR graylags OR leucorodia OR osphronemidae OR bewickii OR "common moorhen" OR "common moorhens" OR decapodiformes OR gobbler OR gobblers OR odontophoridae OR paddlefishes OR salmonine OR esociformes OR "eurasian woodcock" OR "eurasian woodcocks" OR "european smelt" OR "european smelts" OR goldfishes OR tenches OR tyranni OR "common chaffinch" OR "common chaffinchs" OR "common redstart" OR "common redstarts" OR "common roach" OR "common roachs" OR "great knot" OR "great knots" OR potoroidae OR alytidae OR coregonine OR dipteral OR leveret OR "poeciliopsis gracilis" OR amphiumidae OR batrachoidiformes OR "bighead goby" OR heteropneustidae OR lullula OR "norway pout" OR "norway pouts" OR sipunculida OR dogfishes OR sebastidae OR tarsiidae OR alethinophidia OR "common nase" OR "common nases" OR "common sandpiper" OR "common sandpipers" OR "eurasian blackcap" OR "eurasian blackcaps" OR pterocnemia OR syngnathiformes OR "common chaffinches" OR eupleridae OR octopodiformes OR phascolarctidae OR scophthalmidae OR "starry smooth- hound" OR "starry smooth-hounds" OR whitefishes OR cuniculidae OR "european sprat" OR "european sprats" OR "rosy bitterling" OR "rosy bitterlings" OR "common dace" OR "common daces" OR "lesser weever" OR "lesser weevers" OR scaldfish OR "water rail" OR "water rails" OR alouattinae OR centrarchiformes OR "common whitethroat" OR "common whitethroats" OR gavialidae OR "grey gurnard" OR "greygurnards" OR lateolabracidae OR rheiformes OR "tub gurnard" OR "tub gurnards" OR "common chiffchaff" OR "common chiffchaffs" OR garfishes OR "lesser whitethroat" OR "lesser whitethroats" OR myoxidae OR seabasses OR spariformes OR umbridae OR "yellow boxfish" OR anabantiformes OR aotidae OR "common bleak" OR "common bleaks" OR "common rudd" OR "common rudds" OR "greater pipefish" OR hapale OR nandiniidae OR "stone loaches" OR whinchat OR whinchats OR acanthuriformes OR "brotula barbata" OR "common ling" OR "common lings" OR "common roaches" OR cottonrat OR cottonrats OR douroucoulis OR dromaiidae OR fitches OR fitchew OR galaxiiformes OR laprine OR saimiriinae OR solenette OR tarsii OR "tompot blenny" OR "common dragonet" OR "common dragonets" OR "longspinedbullhead" OR "longspined bullheads" OR monotremate OR monotremates OR pempheriformes OR perdicinae OR presbytini OR smegmamorpha OR "bighead gobies" OR "carangaria incertae sedis" OR coiidae OR "fivebeard rockling" OR foulmart OR foumart OR grasskeet OR "greater pipefishes" OR ibices OR millionfish OR muguliformes OR "norwegian topknot" OR peewit OR "red sea sailfin tang" OR rupicapras OR sheatfishes OR "tompot blennies" OR "twait shad" OR "yellow boxfishes")

Results: 10177702

19: #17 AND #18 Results: 87

**\*\*\*\*\*\*\*\*\*\*\*\*\*\*\*\*\*\*\*\*\*\*\*\*\*\*\*\*\*\*\*\*\*\*\*\*\*\*\*\*\*\*\*\*\*\*\*\*\*\*\*\*\*\*\*\*\*\*\*\*\*\*\*\*\*\*\*\*\*\*\*\*\*\*\*\*\*\***

## Appendix 5. Data Extraction Forms

<u>Study Characteristics</u>

1. What is the year of publication?
2. What journal is the article published in?
3. Name of the first author? (Full last name, initial of first name - e.g. Smith, K)
4. Name and email of the corresponding author? (e.g. Smith K)
5. What is the source of funding?

a. Not Reported
b. Government
c. Industry
d. Academic Institution
e. Charity/Foundation
f. Other
g. Unclear
6. What country is the corresponding author from (look at corresponding address)?

<u>Animal Model</u>

1. What is the species?

a. Mouse
b. Rat
c. Monkey
d. Rabbit
e. Other
2. What type of stroke is being studied?

a. Ischemic
b. Hemorrhagic
c. Other
3. What type of stroke model is being used?

a. Intraluminal suture (e.g. Filament, distal, and transient middle cerebral artery occlusion)

i. What is the number of minutes of occlusion time (relevant to lesion severity)?
b. Permanent middle cerebral artery occlusion (i.e. cauterization, permanent clip, permanent distal middle cerebral artery ligation)
c. Photothrombotic
d. Embolism
e. Modified 3 vessel occlusion
f. Endothelin-1
g. Microvascular embolic
h. L-NIO
i. Intracerebral hemorrhagic
j. Other (please specify)
4. What are the sexes of the animals?

a. Male
b. Female
c. Male and female
d. Not reported
5. What is the weight in grams and, if applicable, specify when the weight(s) was measured?
6. What is the age in weeks of the species?
7. Are there comorbidities in the animals have (e.g. aged, obesity, diabetes/hyperglycemia, hypertension)?

a. Yes (please specify)
b. No
8. What specific area of the brain was the stroke induced in?

a. Frontal lobe
b. Parietal lobe
c. Temporal lobe
d. Occipital lobe
e. Cerebellum
f. Thalamus
g. Basal ganglia
h. Other (please describe)
i. Not reported
9. Were there any animals excluded?

a. Yes (please explain why i.e. death, no impairment from stroke, etc.)
b. No
10. What was the total number of animals excluded from the study?

<u>Study Intervention</u>

1. What agent is used?

a. Maraviroc
b. Selzentry
c. Celsentri
d. Leronlimab
e. Aplaviroc
f. Vicriviroc
g. Ancriviroc
h. Other (please specify)
2. What is the dose (numeric value)?
3. What is the unit used for the dose?
4. What is the route of administration?
5. What was the agent prepared/diluted in?
6. When was the treatment administered?

a. Pre-stroke
b. Post-stroke
7. What is the timing in hours of intervention post-stroke (first adminsteration if applicable)?
8. Was the treatment administered at multiple timepoints?

a. Yes (please specify the other timepoints in hours)
b. No
9. How is time zero defined in the study?
10. Was the intervention paired with any other therapy (e.g. post-stroke rehabilitation paradigms)?

a. Yes (please describe)
b. No

<u>Outcomes</u>

1. Are motor behaviour outcomes (e.g. motor skills) being assessed? Please check all that apply

a. Yes

i. Neurological deficit score
ii. Rotarod
iii. Adhesive removal test
iv. Cylinder task
v. Foot fault & paw placement tests (e.g. tapered beam, ladder, grid walking)
vi. Montoya staircase
vii. Reaching task (e.g. tray, pellet, pasta matrix)
viii. Corner test
ix. Staircase test
x. Forelimb placing test
xi. Wire hanging test
xii. Other
b. No
2. Are cognitive behavioural outcomes (e.g. cognitive skills) being assessed? Please check all that apply

a. Yes

i. Morris water maze
ii. ii. Y-maze test
iii. iii. Novel object recognition test
iv. iv. Elevated plus maze
v. v. Sucrose preference test
vi. vi. Tail suspension test
vii. vii. Open field test
viii. viii. Forced swim test
ix. ix. Other
b. No
3. Was infarct volume/size measured?

a. Yes

i. What is the mean infarct size in mm3 or % of the hemisphere for the treatment animals?
ii. What is the mean infarct size in mm3 or % of the hemisphere for the control animals?
iii. What is the method for measuring infarct size? Please check all that apply
  1. Triphenyltetrazolium chloride (TTC)
  2. Cresyl violet (CV)
  3. Hematoxylin and eosin (H&E)
  4. Magnetic resonance imaging (MRI)
  5. Other (please specify)
iv. What is the (first, if applicable) time of the post-stroke measure of infarct size in days?
v. Was infarct size measured multiple times?
  1. Yes (please list the other times in days)
  2. No
b. No
4. How many animals are in the treatment and control groups?
5. What number of animals died for control and treatment groups?
6. What was the adverse effect (e.g. muscle fatigue, reduced mobility) for control and treatment groups
7. Were tissue outcomes (e.g. immunohistochemical staining, axonal neurofilament staining, pharmacogenetic techniques, axonal tracing techniques, genetic markers identification) measured or visualized?

a. Yes, please explain
b. No
8. Were brain imaging or biomarkers used to measure animal neural connectivity?

a. Yes

i. Diffusion tensor imaging (DTI)
ii. ii. Resting state functional MRI (rsFMRI)
iii. iii. Task-based functional MRI
iv. iv. Other (please specify)
b. No

<u>Risk of Bias</u>

<u>“Yes” indicates low risk of bias; “no” indicates high risk of bias; and “unclear” indicates an unclear risk of bias. If one of the relevant signaling questions is answered with “no,” this indicates high risk of bias for that specific entry.</u>

1. Was the allocation sequence adequately generated and applied?

*Did the investigators describe a random component in the sequence generation process such as:

- Referring to a random number table;
- Using a computer random number generator.
  a. Yes
  b. No
  c. Unclear

Additional info:

Examples of a non-random approach:

- Allocation by judgement or by investigator’s preference;
- Allocation based on the results of a laboratory test or a series of tests;
- Allocation by availability of the intervention;
- Sequence generated by odd or even date of birth;
- Sequence generated by some rule based on animal number or cage number.
- 2. Were the groups similar at baseline or were they adjusted for confounders in the analysis?

*Was the distribution of relevant baseline characteristics balanced for the intervention and control groups?

a. Yes
b. No
c. Unclear

*If relevant, did the investigators adequately adjust for unequal distribution of some relevant baseline characteristics in the analysis?

a. Yes
b. No
c. Unclear

*Was the timing of disease induction adequate?

a. Yes
b. No
c. Unclear

Additional info:

The number and type of baseline characteristics are dependent on the review question. Before starting their risk of bias assessment, therefore, reviewers need to discuss which baseline characteristics need to be comparable between the groups. In an SR investigating the effects of hypothermia on infarct size, for example, gender distribution, left ventricular weight and heart rate and blood pressure should be similar between the groups at the start of the study.

A description of baseline characteristics and/or confounders usually contains:

- The sex, age and weight of the animals
- Baseline values of the outcomes which are of interest in the study

Timing of disease induction:

In some prevention studies, the disease is induced after allocation of the intervention. For example, in an experiment on preventive probiotic supplementation in acute pancreatitis, pancreatitis is induced after allocation of the animals to the probiotic or control group. To reduce baseline imbalance, the timing of disease induction should be equal for both treatment groups.

Examples of adequate timing of disease induction:

- The disease was induced before randomization of the intervention.
- The disease was induced after randomization of the intervention, but the timing of disease induction was at random, and the individual inducing the disease was adequately blinded from knowing which intervention each animal received.
- 3. Was the allocation to the different groups adequately concealed during?

*Could the investigator allocating the animals to intervention or control group not foresee assignment due to one of the following or equivalent methods?

- Third-party coding of experimental and control group allocation Central randomization by a third party; Sequentially numbered opaque, sealed envelopes

- Yes
- No
- Unclear

Additional info:

Examples of investigators allocating the animals being possibly able to foresee assignments:

- Open randomization schedule
- Envelopes without appropriate safeguard
- Alternation or rotation
- Allocation based on date of birth
- Allocation based on animal number
- Any other explicitly unconcealed procedure of a non-random approach
- Were the animals randomly housed during the experiment?

*Did the authors randomly place the cages or animals within the animal room/facility?

a. Yes
b. No
c. Unclear

1. Animals were selected at random during outcome assessment (use signaling questions of entry 6).

*Is it unlikely that the outcome or the outcome measurement was influenced by not randomly housing the animals?

a. Yes
b. No
c. Unclear

The animals from the various experimental groups live together in one cage/pasture (e.g., housing conditions are identical).

Additional info:

Examples of investigators using a non-random approach when placing the cages:

- Experimental groups were studied on various locations (e.g., group A in lab A or on shelf A; Group B in Lab B or on shelf B).
- Were the caregivers and/or investigators blinded from knowledge which intervention each animal received during the experiment?

*Was blinding of caregivers and investigators ensured, and was it unlikely that their blinding could have been broken?

a. Yes
b. No
c. Unclear
d. ID cards of individual animals, or cage/animal labels are coded and identical in appearance.
e. Sequentially numbered drug containers are identical in appearance.
f. The circumstances during the intervention are specified and similar in both groups (#).
g. Housing conditions of the animals during the experiment are randomized within the room (use criteria of entry 4).

Additional info:

Examples of inappropriate blinding:

- Colored cage labels (red for group A, yellow group B)
- Expected differences in visible effects between control and experimental groups
- Housing conditions of the animals are not randomized within the room during the experiment; use criteria of entry 4
- The individual who prepares the experiment is the same as the one who conducts and analyses the experiment
- Circumstances during the intervention are not similar in both groups (#)
- Examples where circumstances during the intervention were not similar:
- Timing of administration of the placebo and exp drug was different.
- Instruments used to conduct experiment differ between experimental and control group (e.g., experiment about effects abdominal pressure; exp group receives operation and needle to increase pressure, while control group only has the operation).

**The relevance of the above-mentioned items depends on the experiment. Authors of the review need to judge for themselves which of the above-mentioned items could cause bias in the results when not similar. These should be assessed.

1. Were animals selected at random for outcome assessment?

*Did the investigators randomly pick an animal during outcome assessment, or did they use a random component in the sequence generation for outcome assessment?

a. Yes
b. No
c. Unclear
d. Referring to a random number table;
e. Using a computer random number generator;
f. Etc.

1. Was the outcome assessor blinded?

*Was blinding of the outcome assessor ensured, and was it unlikely that blinding could have been broken?

a. Yes
b. No
c. Unclear
d. Outcome assessment methods were the same in both groups.
e. Animals were selected at random during outcome assessment (use signaling questions of entry 6).

*Was the outcome assessor not blinded, but do review authors judge that the outcome is not likely to be influenced by lack of blinding?

a. Yes
b. No
c. Unclear

(e.g., mortality)

Additional info:

This item needs to be assessed for each main outcome.

1. Were incomplete outcome data adequately addressed? (*)

*Were all animals included in the analysis?

a. Yes
b. No
c. Unclear

*Were the reasons for missing outcome data unlikely to be related to true outcome? (e.g., technical failure)

a. Yes
b. No
c. Unclear

*Are missing outcome data balanced in numbers across intervention groups, with similar reasons for missing data across groups?

a. Yes
b. No
c. Unclear

*Are missing outcome data imputed using appropriate methods?

a. Yes
b. No
c. Unclear

1. Are reports of the study free of selective outcome reporting? (*)

*Was the study protocol available and were all of the study’s pre-specified primary and secondary outcomes reported in the current manuscript?

a. Yes
b. No
c. Unclear

*Was the study protocol not available, but was it clear that the published report included all expected outcomes (i.e. comparing methods and results section)?

a. Yes
b. No
c. Unclear

Additional info:

Selective outcome reporting:

- Not all of the study’s pre-specified primary outcomes have been reported;
- One or more primary outcomes have been reported using measurements, analysis methods or data subsets (e.g., subscales) that were not pre-specified in the protocol;
- One or more reported primary outcomes were not pre-specified (unless clear justification for their reporting has been provided, such as an unexpected adverse effect);
- The study report fails to include results for a key outcome that would be expected to have been reported for such a study.
10. Was the study apparently free of other problems that could result in high risk of bias? (*)

*Was the study free of contamination (pooling drugs)?

a. Yes
b. No
c. Unclear

*Was the study free of inappropriate influence of funders?

a. Yes
b. No
c. Unclear

*Was the study free of unit of analysis errors?

a. Yes
b. No
c. Unclear

*Were design-specific risks of bias absent?

a. Yes
b. No
c. Unclear

*Were new animals added to the control and experimental groups to replace drop-outs from the original population?

a. Yes
b. No
c. Unclear

Additional info:

The relevance of the signaling questions (Table 3) depends on the experiment. Review authors need to judge for themselves which of the items could cause bias in their results and should be assessed.

Contamination/pooling drugs:

Experiments in which animals receive besides the intervention drug additional treatment or drugs which might influence or bias the result.

Unit of analysis errors:

- Interventions to parts of the body within one participant (i. e., one eye exp; one eye control).
- All animals receiving the same intervention are caged together, but analysis was conducted as if every single animal was one experimental unit.
- Design-specific risks of bias:
- Crossover design that was not suitable (intervention with no temporary effect, or the disease is not stable over time)
- Crossover design with risk of carry-over effect
- Crossover design with only first period data being available
- Crossover design with many animals not receiving 2nd or following treatment due to large number of drop-outs probably due to longer duration of study
- Crossover design in which all animals received same order of interventions
- Multi-arm study in which the same comparisons of groups are not reported for all outcomes (selective outcome reporting)
- Multi-arm study in which results of different arms are combined (all data should be presented per group)
- Cluster randomized trial not taking clustering into account during statistical analysis (unit of analysis error)
- Crossover design in which paired analysis of the results is not taken into account

### Validity

1. Does the treatment effect vary with dose?

a. Yes
b. No
c. Not applicable (only tested one dose)
d. Unsure
2. Does the treatment remain effective when administered at clinically relevant delayed times?

a. Yes (tested after 6 hours and the CCR5 antagonist was still effective)
b. No (tested after 6 hours, but the CCR5 antagonist was not effective)
c. Not applicable (not tested after 6 hours)
d. Unsure
3. Does the treatment cause expected physiological effects?

a. Yes
b. No
c. Unsure
4. Does the treatment penetrate the blood brain barrier?

a. Yes
b. No
c. Unsure
5. What the tests done across multiple laboratories?

a. Yes
b. No
c. Unsure
6. Was testing done on gyrencephalic species (non-human primates i.e. monkeys, dogs, pigs, etc.)

a. Yes
b. No
c. Unsure
7. Did the preclinical testing of therapy occur during the awake phase for the animal model (during the dark phase for rodents i.e. was the test done during the nighttime)?

a. Yes
b. No
c. Unsure

**\*\*\*\*\*\*\*\*\*\*\*\*\*\*\*\*\*\*\*\*\*\*\*\*\*\*\*\*\*\*\*\*\*\*\*\*\*\*\*\*\*\*\*\*\*\*\*\*\*\*\*\*\*\*\*\*\*\*\*\*\*\*\*\*\*\*\*\*\*\*\*\*\*\*\*\*\*\***

## Notes

https://doi.org/10.7554/eLife.103245.2

### Competing Interest Statement

The authors have declared no competing interest.

### Summary of Updates

Updated main text throughout the manuscript to address reviewer and editorial comments received from eLife review.

